# Tail length of triazine-based lipids influences blood clotting risk in vitro and in vivo

**DOI:** 10.64898/2026.08.25.747137

**Authors:** Nabilah Ibnat, Abdullah Al Masud, Julian Mory, Timothy Funk, Dlovan F D Mahmood, Jeremy P Wood, Vincent J Venditto

## Abstract

Lung-targeted delivery of mRNA with lipid nanoparticles (LNPs) demonstrates high potential for therapeutic applications in pulmonary disorders. However, progress in pulmonary mRNA therapeutics is constrained by the challenges of engineering lipids that are both safe and highly effective at targeting the lungs. To meet these critical needs, we designed triazine-based (TZ) ionizable lipids with cyanuric chloride as the linker between the cationic head and the lipophilic tail, which allows for easy derivatization capable of systemic mRNA delivery. Three TZ-based lipids were synthesized using the same ionizable headgroups while differing in the carbon tail length and evaluated for their in vitro and in vivo protein expression. Notably, all three lipids result in pulmonary expression after intravenous administration, but the TZ lipid containing a C14 tail does so without any indication of thrombosis, both in vitro and in vivo as compared to other formulations. Our findings highlight the effect of minor chemical modifications driving altered in vivo activity, thus enabling new opportunities for safe pulmonary delivery of mRNA for lung-related diseases.

**Highlights:**

- Triazine-based ionizable lipids deliver mRNA to extrahepatic tissues
- Tail length of TZ lipid modulates thrombosis potential in the lung
- TZ lipid with C14 tail length exhibits minimal to no observed clotting in mice and human plasma

**GRAPHICAL ABSTRACT:** 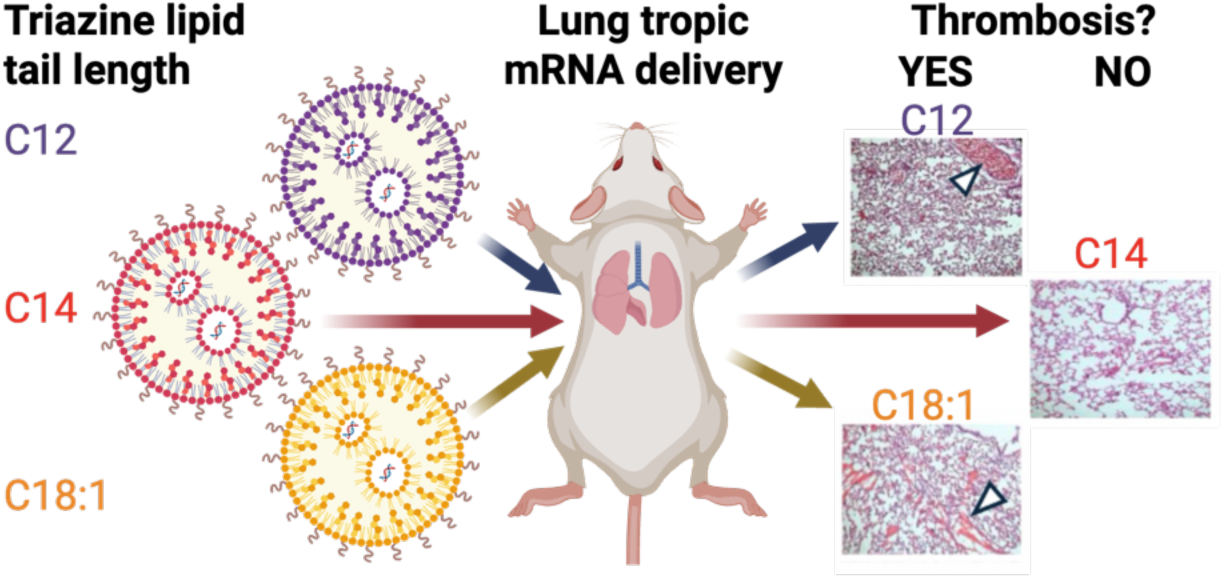

## 1. Introduction

With the prominent progress of mRNA vaccines and the approval of siRNA therapeutics by the FDA, lipid nanoparticles (LNPs) have become the most clinically advanced delivery vehicles for nucleic acid therapeutics.^1–4^ Despite recent advances, nanotherapeutics tend to accumulate in the liver and spleen following systemic delivery, due to lipid composition, and undergo uptake by the phagocyte mononuclear system. ^5,6^ This rapid clearance of mRNA-LNP therapeutics by the liver and spleen limits their biodistribution to other vital organs^7^, such as the lung, kidney, brain, and retina, and highlights the need for further exploration delivery vehicles for extrahepatic mRNA delivery. Targeted mRNA delivery to such organs enables the expression of therapeutic proteins to treat various genetic diseases, cancers, or acute injuries.^8,9^ For successful mRNA expression in extrahepatic organs, mRNA-LNPs encounter multiple barriers, including precise targeting and cellular internalization in specific cell types.^10^ To achieve organ-specific mRNA delivery, efforts are underway to investigate active and passive targeting strategies. Active targeting involves the use of ligands decorated on the LNP surface, such as peptides, proteins, antibodies, and small molecules, to direct LNPs to specific cell receptors.^11–14^ Conversely, passive targeting relies on the design of LNP by adjusting lipid composition and ratio to enhance the development of endogenous protein corona and delivery to specific cells.^15^

Typically, the route of administration affects mRNA delivery, expression and the resultant in vivo activity.^16^ Although for respiratory diseases, local administration (e.g., inhalation) can be efficient, systemic delivery (e.g., intravenous) of the mRNA payload is preferred to achieve rapid and concentrated drug delivery with reproducible pharmacokinetics and pharmacodynamic responses.^17^ Recently, for lung-targeted mRNA delivery via the systemic route, several LNP systems have been developed.^18–20^ A recent study demonstrated that lung-targeted LNPs can be fabricated by incorporating a cationic lipid (e.g. DOTAP) into the four-component system to direct lung tropism after systemic injection.^10^ Similarly, replacement of the helper phospholipid (DOPE) with a permanently charged cationic lipid also facilitated targeted mRNA delivery to the lung epithelial cells.^21^ These findings reinvigorated efforts to modify LNP formulations to achieve desirable delivery profiles and have launched a new wave of discoveries for extrahepatic LNP design.

Systemic administration of lung-targeted systems offers the potential to treat acute and chronic pulmonary conditions with dose-dependent responses and rapid onset of action. In particular, extensive work using lipids containing different cationic amine head groups for pulmonary delivery has been reported,^22–24^ which has expanded to cationic siloxane-containing lipids,^25^ and sulfonium-containing cationic head groups.^26^ The hydrophobic tail region of ionizable lipids also determines organ specificity and biodistribution.^27,28^ Several tail modifications have been investigated through the introduction of an amide bond in the hydrocarbon tail,^29^ as well as branched-tail LNPs for systemic mRNA delivery to the lung.^24,30^ Despite these advances, a thorough assessment of toxicity arising from the systemic administration of cationic nanoparticles in mouse models remains lacking. Importantly, successful protein expression in the lungs has been correlated with the risk of cationic lipids to induce pulmonary thrombosis.^31^ Due to the specialized and vascularized structure of lung tissues, it is essential to carefully evaluate the blood clotting properties of the mRNA-LNP formulations targeting lung tissues for protein expression to ensure their safety.

Previously, we reported the synthesis of a library of triazine lipids through the incorporation of commercially available small molecule head groups and various dialkyl amines as lipid tails.^32^ The triazine (TZ) lipids are efficiently synthesized using cyanuric chloride as a linker and represent a novel class of lipids capable of delivering nucleic acids both in vitro and in vivo.^33^ The objective of this study was to assess the mRNA delivery and protein expression profile of three candidate TZ lipids containing the same head groups and different lipid tails (C12, C14, C18:1) for in vitro and in vivo applications. While all three lipids exhibit robust protein expression in vitro and in vivo, the unexpected observation of tail-dependent pulmonary thrombosis suggests new avenues to modify and mitigate extrahepatic tropism and toxicity through minor chemical modifications. These data serve as a platform to launch additional efforts in the rational design of lipids for safe and effective delivery of mRNA to lung tissue.

## 2. Materials and Methods

### 2.1 Materials

DSPC (1,2-distearoyl-sn-glycero-3-phosphocholine), DOPE (1,2-dioleoyl-sn-glycero-3-phosphoethanolamine), DOTAP (1,2-dioleoyl-3-trimethylammonium-propane (chloride salt)), DSPE-PEG2000 (1,2-distearoyl-sn-glycero-3-phosphoethanolamine-N-(polyethylene glycol)-2000, DMG-PEG2000 (1,2-dimyristoyl-rac-glycero-3-methoxypolyethylene glycol-2000), and cholesterol were purchased from Avanti Polar Lipids, Inc. ALC-0315 was procured from Sinopeg, and cKK-E12 was purchased from Echelon Biosciences. DMEM culture medium, fetal bovine serum (FBS), and sodium acetate buffer solution were procured from Thermo Fisher Scientific. The TNS assay reagents were purchased from Sigma Aldrich. The cell viability kit for MTS assay was obtained from Promega, and the Quant-iT RiboGreen RNA reagent was procured from Invitrogen. Firefly Luciferase (FLuc) and Green Fluorescent Protein (GFP) mRNA were obtained from Trilink BioTechnology. Powder luciferin reagent (for in vivo assay) was from Regis Technologies and D-luciferin free acid (for in vitro assay) was procured from Nanolight to prepare luciferin reagent for luciferase assay, using a previously reported protocol.^34,35^ Amicon Ultra Centrifugal Filters (10 kDa MWCO) were purchased from MilliPore Sigma. The thrombin-antithrombin (TAT) complex ELISA kit and H&E staining kit were obtained from Abcam.

### 2.2 Methods

#### Synthesis, purification, and analysis of triazine lipids

Triazine lipids were synthesized as previously reported.^32^ Briefly, cyanuric chloride was reacted with nucleophilic lipid tails under an inert atmosphere at 4 °C for 1.5 h in basic conditions to yield the dichlorotriazine. The second and third nucleophilic aromatic substitutions proceeded with mono-protected diamine headgroups under inert atmosphere in basic conditions at 45 °C for 20 h, then 82 °C for 20 h. The resulting compounds were isolated and treated with trifluoroacetic acid to afford the final lipid products. All ^1^H and ^13^C{^1^H} NMR spectra were recorded at ambient temperature at 400 MHz and 100 MHz, respectively, on a Bruker Avance Neo 400 MHz FT-NMR spectrometer unless otherwise noted. Chemical shifts are reported in parts per million (ppm) relative to tetramethylsilane (TMS) for spectra taken in CDCl_3_. Samples were prepared in 0.7 mL of solvent unless otherwise noted. High-resolution mass spectrometry data were collected at the Johns Hopkins University Mass Spectrometry Facility. Analytical thin-layer chromatography (TLC) was performed using silica gel 60 F254 precoated plates (0.25 mm thickness) with a fluorescent indicator. Flash column chromatography was performed using silica gel 60 (230-400 mesh).

#### Formulation and Characterization of Triazine Lipid Nanoparticles (TZ-LNPs)

To prepare TZ-LNPs, triazine lipids were mixed with DOPE, cholesterol, and DSPE-PEG 2000 at a molar ratio of 50/10/38.5/1.5 in ethanol. For formulations containing ALC-0315, the ethanol mixture was prepared using the commercial ionizable lipid ALC-0315, DSPC, cholesterol and DMA-PEG 2000 at a molar ratio of 46/9.4/42.9/1.7, as previously reported.^36^ DOTAP formulations consisted of DOTAP, cKK-E12, DOPE, cholesterol, and DMG-PEG 2000 at a molar ratio of 50/25/5/18.5/1.5, as previously reported.^31^ The mRNA was diluted in acetate buffer (25 mM, pH 4.0) to prepare the aqueous phase. LNPs were formulated with mRNA using a microfluidic mixing method (NanoAssemblr Ignite, Precision Nanosystems (now Cytiva)) at an aqueous: organic flow-rate ratio (FRR) of 3:1 and a total flow rate (TFR) of 12 mL/min. The total lipid/mRNA weight ratio was 40:1. Following formulation, the mRNA-LNPs were diluted in sterile sucrose buffer (254 mM) with 10.7 mM sodium acetate and 20 mM Tris-buffer (pH 7.4). Ethanol removal was performed using Amicon Ultra Centrifugal Filter (10KDa MWCO), at 4 °C, 3500 x g centrifugation speed. The concentrated mRNA-LNPs were then used for subsequent in-vitro and in-vivo experiments. Hydrodynamic sizes, polydispersity Index (PDI) and zeta potential (ZP) of mRNA-LNPs were measured by Malvern Zetasizer Nano-ZS using dynamic light scattering (DLS) technology. Samples were mixed in sucrose buffer at 1:20 ratio and tested in DTS 1070-folded capillary cells at 25 mV, and data were analyzed using ZS Xplorer software.

#### Ribogreen Assay for mRNA Encapsulation Efficiency

The Quant-it Ribogreen assay was used to quantify mRNA concentration and encapsulation efficiency of the mRNA-LNP formulations. Briefly, serially diluted mRNA samples for a 6-point standard curve ranged from 250 ng/µL to 10 ng/µL, and mRNA-LNP samples at a target concentration of 100 ng/µL mRNA were prepared in TE buffer. Samples were incubated for 10 min at 37°C in a black 96-well microplate, both in the presence or absence of 0.2% (v/v) Triton X-100/TE to disrupt the LNPs. Following the addition of Ribogreen reagent, fluorescence intensity (Ex: 485 nm/Em: 528 nm, gain: 80) was measured using Biotek 5 plate reader. The following formula was used to calculate mRNA encapsulation efficiency: EE% = [(total mRNA-free mRNA)/total mRNA] X 100%.

#### TNS Assay for *pK_a_*

LNP p*K*_a_ was determined using the TNS binding assay as previously reported.^37^ Briefly, a master buffer stock was prepared containing 7.5 mM sodium acetate, 15 mM sodium borate, 20 mM sodium chloride, 7.5 mM glycineglycine, and 7.5 mM imidazole. Buffers were titrated with HCl or NaOH to achieve pH values between 3 and 10. The 6-(*p*-toluidino)-2-naphthalenesulfonic acid sodium salt (TNS) was prepared as a 300 μM stock solution in DMSO. Based on Ribogreen mRNA concentration, the mRNA-LNPs were diluted in water to a 24.2 µM ionizable lipid concentration and tested in a 96-well black assay plate using the TNS buffer at each pH buffer. The fluorescence intensity (Ex: 321 nm/Em: 445 nm, gain: 60) of the test samples was measured using Biotek 5 plate reader. Data were fitted with a sigmoidal curve of fluorescence intensity vs. concentration, and p*K*_a_ was calculated as a log transformation from the inflection point of the curve.

#### In vitro mRNA Transfection

For in vitro screening, a cell density of 50,000 HEK293T cells/well was seeded in a clear-bottom 24-well plate and grown to 70-80% confluency for 24 h at 37 °C under 5% CO_2_ atmosphere. The cells were treated with EGFP mRNA-loaded LNPs with 500 ng mRNA in triplicate and were incubated for an additional 24 hr. Then, cells were detached using Trypsin and resuspended in PBS buffer containing 0.1% BSA and 1 mM EDTA to make a single-cell suspension. Finally, flow cytometry was performed using a 488-nm excitation laser, measuring 10,000+ fluorescence events including forward scatter and side scatter, to quantify the percentage of EGFP-positive cells. Cells were identified as positive based on enhanced green fluorescence intensity compared to the negative control.^38^

#### In Vitro Cytotoxicity Assay

HEK293T cells were cultured in DMEM medium containing 10% FBS and 1% streptomycin and penicillin at 37 °C under 5% CO_2_ atmosphere. For testing in vitro cell viability, the HEK293T cells were seeded into 96-well plates at a density of 25,000 cells per well and incubated at 37 °C under 5% CO_2_ atmosphere overnight. The next day, the cells were treated with different concentrations of lipids ranging from 12.5 µM to 200 µM. After 4 h of incubation, the cell media were replaced with fresh DMEM and incubated for another 24 h. Then, relative cell viability was examined by MTS reagent according to the manufacturer’s instructions (Promega).

#### Blood Collection and Calibrated Automated Thrombogram (CAT) Assay

Plasma thrombin generation in human plasma was measured using the CAT assay as previously described.^39,40^ Human pooled normal plasma from CRYOcheck (CCN10-10) was used. In brief, frozen plasma samples were thawed for 10 min at 37 °C and incubated with LNPs containing 500 ng of mRNA for 5 min. In a 96-well plate, the mRNA-LNP-treated plasma samples were divided into two subsamples; the first subsample was without both phospholipid (PL) and kaolin as a coagulator, and the second subsample contained both PL (4 µM, consisting of 20% phosphatidylserine, 20% phosphatidylethanolamine, 60% phosphatidylcholine) and kaolin (10 µg/mL). Thrombin generation was measured following addition of a fluorogenic substrate for thrombin (Z-Gly-Gly-Arg-AMC) plus calcium chloride. The fluorescent measurements were obtained continuously for up to 2 hours by fluorimeter. The thrombin measured over time was calculated using Thrombinoscope software and derived parameters included: lag time (initiation phase of coagulation); endogenous thrombin potential (ETP; area under curve); peak height (PH; maximum level of thrombin generated); and time to peak.

#### In Vivo Studies

##### Mice

All animal experiments were approved by the Institutional Animal Care and Use Committee (IACUC) of the University of Kentucky (protocol 2020-3523). Male and female Balb/c mice (6-8 weeks old) were purchased from the Jackson Laboratory. All mice were housed in a pathogen-free facility and handled according to the animal welfare guidelines of the University of Kentucky.

##### In vivo Biodistribution Protein Expression

For FLuc mRNA delivery and bioluminescence imaging, Balb/c mice were sedated with isoflurane and administered PBS or mRNA-LNPs (0.25 mg mRNA/kg body weight) through retro-orbital injection. At 4 h after treatment, 150 µl of D-luciferin in sterile PBS was injected intraperitoneally. After 10 min, bioluminescence imaging was performed on the sedated mice using Spectral Instruments Imaging Ami-HT system. Mice were then sacrificed, and internal organs (liver, lung, spleen, kidney and heart) were collected for ex-vivo imaging. The signal intensity was normalized by subtracting the background signal and analyzed using Aura (Ami-HT) software to evaluate the FLuc protein expression in each organ.

##### Lung Histology

For histological analysis, Balb/c mice were administered PBS or mRNA-LNPs (0.25 mg mRNA/kg body weight) by retro-orbital injections. After 30 min, mice were sacrificed, the abdominal cavity was cut open, and the heart was perfused with 5 mL PBS. Next, the lungs were inflated with 10% buffered formalin (BF) and collected in formalin solution for overnight fixation according to a published protocol.^41^ The next day, the lung tissues were transferred to 20% sucrose solution for 24-48 hours and then embedded in OCT blocks for storage at -80 °C. Within one month, the OCT-embedded lung tissues were sectioned at 5-µm-thickness using a cryostat and stained with hematoxylin and eosin (H&E) according to the manufacturer’s instructions (Abcam). The stained slides were imaged using an Echo Revolve Microscope.

##### Thrombin-Anti-Thrombin (TAT) Assay

The mRNA-LNP formulations were injected into Balb/c mice (n=6) for a circulation time of 30 minutes. Blood was collected from mice into K_2_EDTA tubes by submandibular venipuncture. The tubes were immediately centrifuged at 1500 x g for 10 min. Plasma was collected and analyzed with a TAT-ELISA kit (Abcam) according to the manufacturer’s instructions.

##### Statistical analysis

Data are presented as mean ± standard deviation. All statistical analyses were carried out using GraphPad Prism 10 (GraphPad Software Inc.). All experiments were carried out in triplicate, and statistical comparisons were performed using one-way analysis of variance (ANOVA) for in vivo analysis, and statistical significance indicated as \**p* < 0.05, \*\**p* < 0.01, and \*\*\**p* < 0.001.

## 3. Results

### Lipid synthesis

Triazine lipids were synthesized using cyanuric chloride as a linker between alkyl amine tails and diamine headgroups, as previously reported.^32^ Cyanuric chloride is an ideal linker for lipid synthesis as it undergoes thermally controlled nucleophilic aromatic substitution (S_N_Ar) at each electrophilic carbon to efficiently yield diverse lipid molecules. The three lipids described herein are based on previous efforts using the C12 derivative for plasmid DNA delivery,^33^ and our efforts to investigate the effect of tail length on nucleic acid delivery. Therefore, three lipids were synthesized using one of three dialkylamine nucleophiles (C12 – didodecylamine, Scheme S1; C14 – ditetradecylamine, Scheme S2; C18:1 – dioleylamine, Scheme S3) to afford a dichlorotriazine substituted with lipid tail. All three dichlorotriazines were then reacted with two equivalents of mono-Boc-protected diaminopropane and subsequently deprotected under acidic conditions to yield each of the three ionizable lipids containing the same headgroups and three different lipid tails. The three-step lipid syntheses proceeded in 34-45% overall yield.

### Physicochemical characterization and in vitro protein expression of TZ-LNPs

LNPs containing each of the newly synthesized lipids (C12, C14, C18:1) were formulated with firefly luciferase (FLuc) mRNA using a molar composition of 50:10:1.5:38.5 (TZ lipid:DOPE:DSPE-PEG2000:Chol) at a weight ratio of 40:1 for lipid to mRNA. Control formulations containing ALC-0315 and DOTAP were prepared using optimized compositions previously reported in the literature **(Table S1).** Physicochemical properties of the resulting formulations were characterized to assess differences in nanoparticle critical quality attributes. The size of the formulation containing DOTAP was statistically larger as compared to the size of formulations containing either ALC-0315 or the TZ lipids (**Figure 1A**). The mean diameters of ALC-0315 and DOTAP formulations were 101.6 ± 10.5 nm and 156.0 ± 23.7 nm, respectively. The diameters of mRNA-LNP formulations containing C12, C14 and C18:1 tails were 101.9 ± 10.3 nm, 94.7 ± 3.4 nm, and 87.4 ± 8.3 nm, respectively. Additionally, the polydispersity index (PDI) for all formulations was between 0.2 and 0.3, indicating reasonably uniform size distributions (**Figure 1A**). The zeta potential (ZP) values for ALC-0315 and three TZ lipid formulations were within -10 mV to +10 mV, while DOTAP formulations exhibited a slightly positive ZP value of 15.5 ± 3.9 mV (**Figure 1B**). Notably, the mRNA encapsulation efficiency of all three TZ-LNPs and DOTAP were more than 90%, and significantly higher than that of formulations prepared with ALC-0315 LNPs (62.9 ± 15.3%, \*\**p* < 0.01; **Figure 1C**). The p*K*_a_ of each LNP was then assessed by TNS assay **(Table S2)**, resulting in values of 8.85 ± 7.38, 7.07 ± 0.91 and 9.0 ± 1.63 for C12, C14 and C18:1 LNPs, respectively. Notably, all three TZ lipids exhibit reduced fluorescence magnitude and demonstrate more gradual curves lacking a sharp transition observed in ALC-0315 (p*K*_a_ = 6.41 ± 0.28), which is consistent with previous reports.^42,43^ This is most pronounced with C12 TZ LNP, which lacks a clear transition with a confidence interval beyond the limits of the range tested.

**Figure 1:**
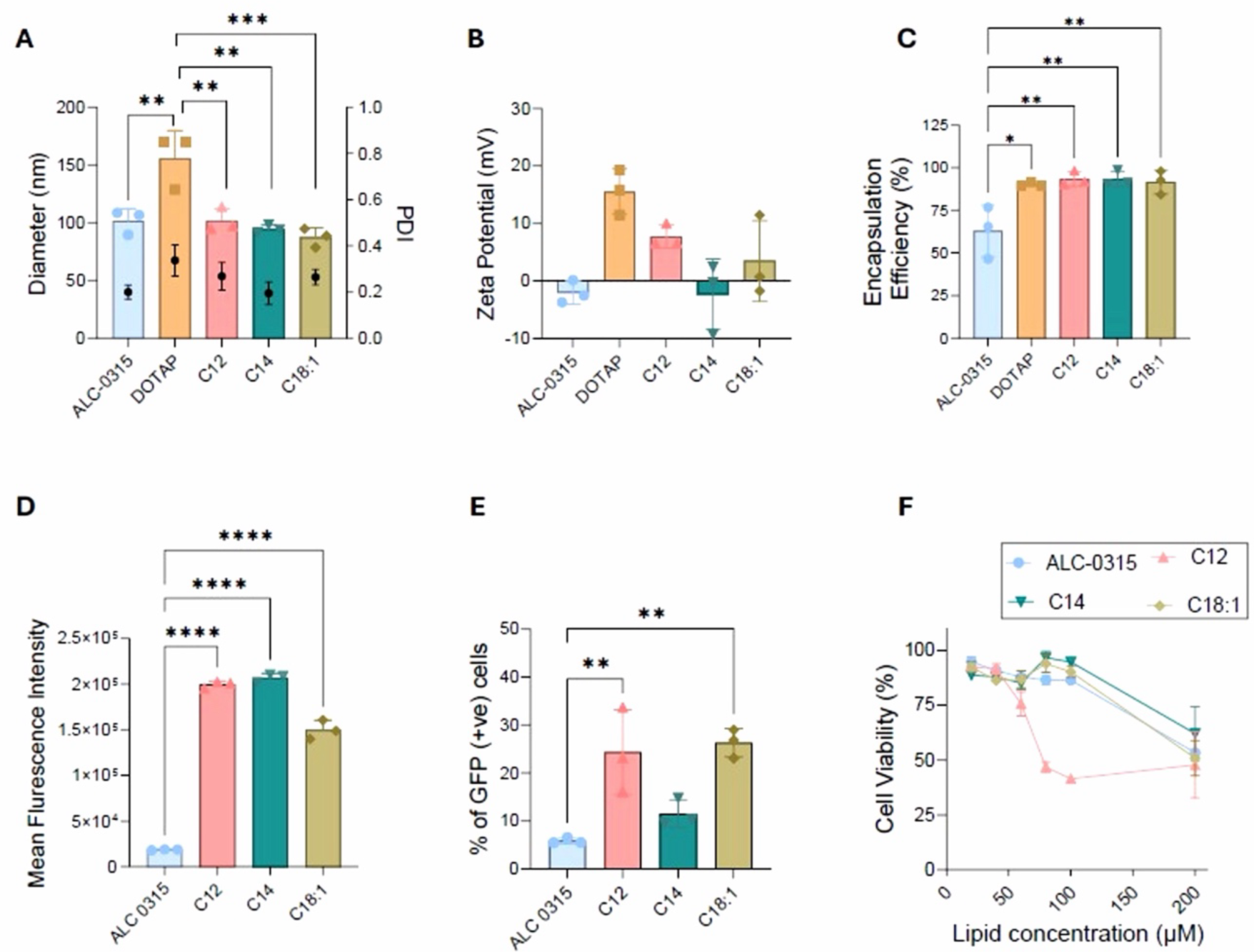
Physicochemial characterization of FLuc mRNA-LNP formulations prepared with C12, C14, C18:1 triazine lipids using microfluidic mixing method at a flow rate of 12 mL/min. (A) Diameter of particles (nm) with PDI, (B) Zeta Potential, (C) mRNA encapsulation efficiency (EE). The size, PDI and ZP were measured by DLS, and EE was measured using the Ribogreen assay. In vitro efficiency in HEK-293T cells transfected with EGFP mRNA-loaded TZ-LNPs by flow cytometry. (D) Mean Fluorescence Intensity, (E) Percentage of GFP-positive cells. (F) Cell viability of TZ lipids with increasing lipid concentrations (12.5 μΜ το 200 μM) in HEK-293T cells using MTS assay. For each set of experiments, n=3 replicates are shown. Statistical significance is indicated as follows: (*) *p*<0.05, (**) *p*<0.01, (***) *p*<0.001, (****) *p*<0.0001.

To assess mRNA delivery and protein expression in vitro, cell studies were performed using each of the three triazine lipids. Protein translation was quantified in HEK293T cells using flow cytometry after delivery of EGFP mRNA as a reporter protein. All three TZ-LNPs exhibit statistically increased intracellular mean fluorescent intensity (MFI) as compared to formulations containing ALC-0315 (**Figure 1D, \****p*<0.01). The MFI for ALC-0315, C12, C14, and C18:1 was (0.2 ± 0.0) x 10^5^; (1.9 ± 0.04) x 10^5^; (2.0 ± 0.05) x 10^5^; and (1.4 ± 0.1) x 10^5^, respectively. Additionally, the percentage of GFP-positive cells was significantly increased when delivering mRNA with either C12 (24.3 ± 8.0%) or C18:1 (26.3 ± 3%) lipids as compared to ALC-0315 (5.9 ± 0.6%, \*\**p*<0.001) (**Figure 1E**). These data indicate that TZ-LNPs provide more efficient GFP expression in vitro as compared to ALC-0315.

The cytotoxicity of each TZ-LNP was also investigated in HEK293T cells by MTS assay. The cell viability data at different lipid concentrations indicate that C14 and C18:1 TZ lipids exhibit >90% cell viability at lipid concentrations 100 µM and below, similar to LNPs prepared with ALC-0315. However, the C12 TZ lipid exhibits decreased cell viability at concentrations greater than 50 µM (**Figure 1F**).

### Evaluation of in vivo FLuc mRNA expression using TZ-LNPs

To explore the in vivo mRNA delivery properties of TZ-LNPs, 0.25 mg/kg dose of firefly luciferase (FLuc) mRNA was formulated and intravenously administered to Balb/c mice. At 4 hr post-treatment, bioluminescence images of the whole mouse and ex vivo organs were collected. Image analysis indicates that the three TZ-based LNPs (C12, C14, C18:1) promoted prominent FLuc expression in the lungs as compared to hepatic expression in the liver when using ALC-0315. Protein expression was not observed in the heart and kidney for any of the formulations tested (**Figure 2A and S1**). The total bioluminescence emission flux in the lungs of mice receiving PBS showed a mean signal intensity of (4.8 ± 2.3) x 10^3^ p/s. Formulations containing DOTAP exhibited more than 6 times greater flux in the lungs (1.8 ± 0.7) x 10^8^ p/s as compared to ALC-0315 (2.8 ± 0.4) x 10^7^ p/s. The TZ-based FLuc mRNA formulations with C12, C14, and C18:1 tail exhibited lung signal intensity of (1.3 ± 0.4) x 10^6^ p/s; (4.9 ± 1.4) x 10^6^ p/s and (3.4 ± 1.8) x 10^6^ p/s, respectively, which was significantly lower than the DOTAP formulation; but similar to ALC-0315 (**Figure 2B**). Conversely, the signal intensity in the liver was higher with ALC-0315 (4.4 ± 2.3) x 10^9^ p/s as compared to the DOTAP (8.2 ± 6.3) x 10^6^ p/s formulation, while all three TZ formulations C12, C14, and C18:1 exhibit signal intensity of (1.1 ± 0.7) x 10^6^ p/s; (0.1 ± 0.04) x 10^6^ p/s and (0.1 ± 0.08) x 10^6^ p/s, respectively (**Figure 2C**). The signal intensity in the lung and liver was then normalized to tissue weight and a lung/liver ratio was calculated for each mouse, indicating a 20-fold increase in protein expression in the liver as compared to the lung for ALC-0315, while DOTAP exhibits a greater than 100-fold increase in lung expression as compared to liver. All three TZ lipids exhibit similar expression in the lung, with the C12 lipid exhibiting 10-fold increase in lung over liver, and C14 and C18:1 lipids exhibiting more than a 100-fold increased translation in the lung over liver (**Figure 2D**). Notably, the C14 and C18:1 TZ lipids also exhibit significantly increased expression in the spleen as compared to DOTAP and ALC-0315, which was not observed with the C12 lipid (**Figure 2E**).

**Figure 2:**
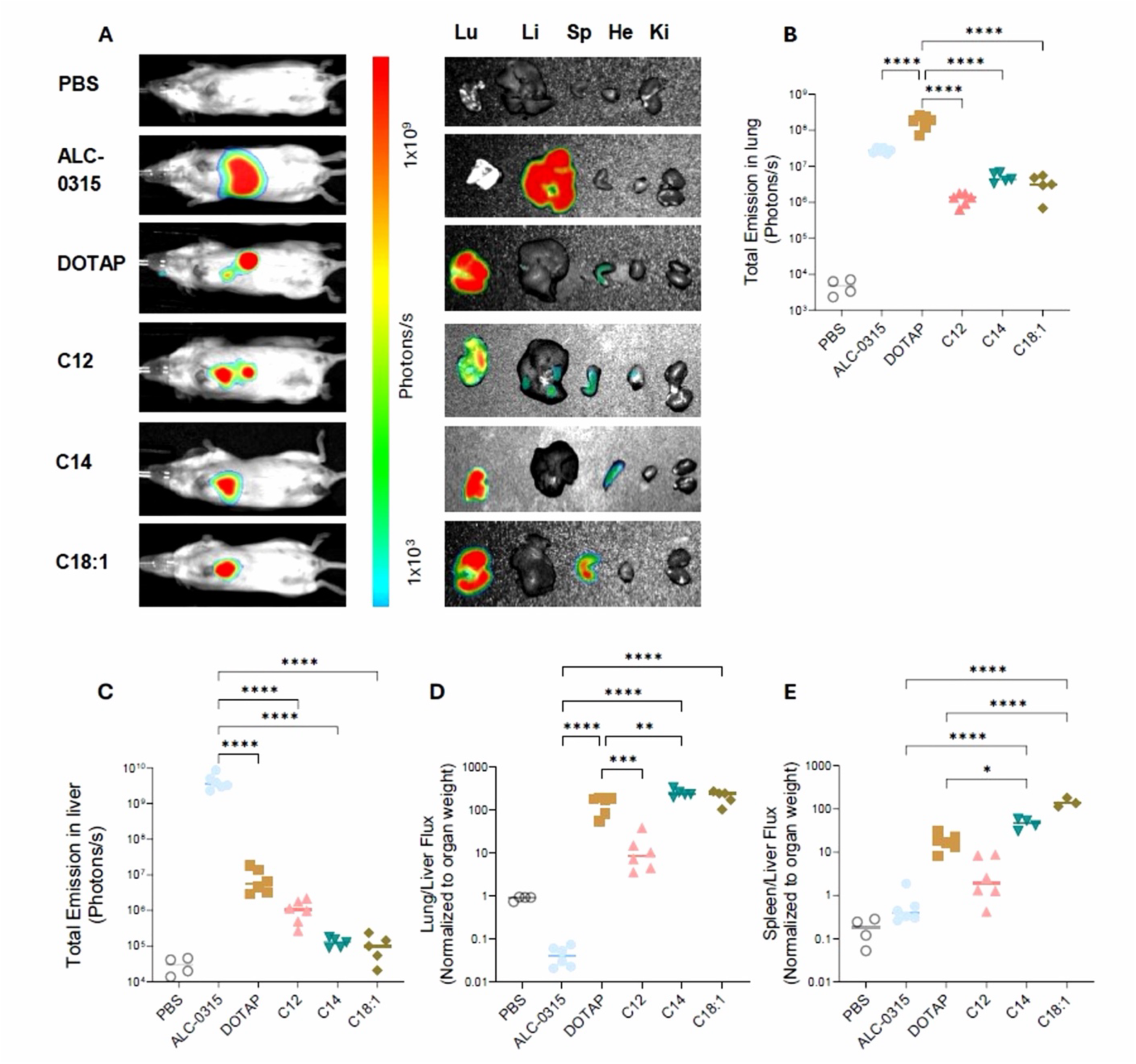
In vivo biodistribution of FLuc protein expression in mice after intravenous administration of LNP-mRNA formulations prepared with C12, C14, C18:1 TZ lipids. (A) Representative whole-body image of mice treated with FLuc mRNA dose of 0.25 mg/kg per mouse; with ex vivo bioluminescence images of major organs (lung, liver, spleen, heart, and kidney) (n=6). PBS was used as a control. (B) Quantification of total flux intensity in the lung, (C) total flux intensity in the liver, (D) total flux intensity ratio in lung/liver, and (E) total flux intensity ration in spleen/liver normalized to organ weight. Statistical significance is indicated as follows: (*) *p*<0.05, (**) *p*<0.01, (***) *p*<0.001, (****) *p*<0.0001.

### Lung histology for blood clotting assessment of TZ-LNPs

To evaluate the blood clotting properties of the TZ-LNPs in lung tissue, a second set of Balb/c mice received 0.25 mg/kg dose of firefly luciferase (FLuc) mRNA formulated in TZ-LNPs via intravenous injection. Three mice were treated in each treatment group with PBS as a negative control. Excised lung tissues collected at 4 h after LNP administration were sectioned and stained with hematoxylin-eosin (H&E) stain to assess the extent of thrombosis. Notably, lung tissues from mice receiving either C12 and C18:1 TZ-LNPs displayed visible blood clots similar to those observed in mice receiving DOTAP formulations (**Figure 3**). On the other hand, mice receiving the C14 TZ-LNP had minimal blood clots and lung morphology similar to mice treated with ALC-0315 and the PBS control (**Figure 3**). Lung histology images (4X magnification) for all mouse groups in triplicate are provided in **Figure S2**. These data suggest that the lipid tail length is a significant driver of the thrombosis potential in mice after systemic administration, with LNPs formed with the C14 TZ-lipid exhibiting minimal clotting as compared to the C12 and C18:1 formulations.

**Figure 3:**
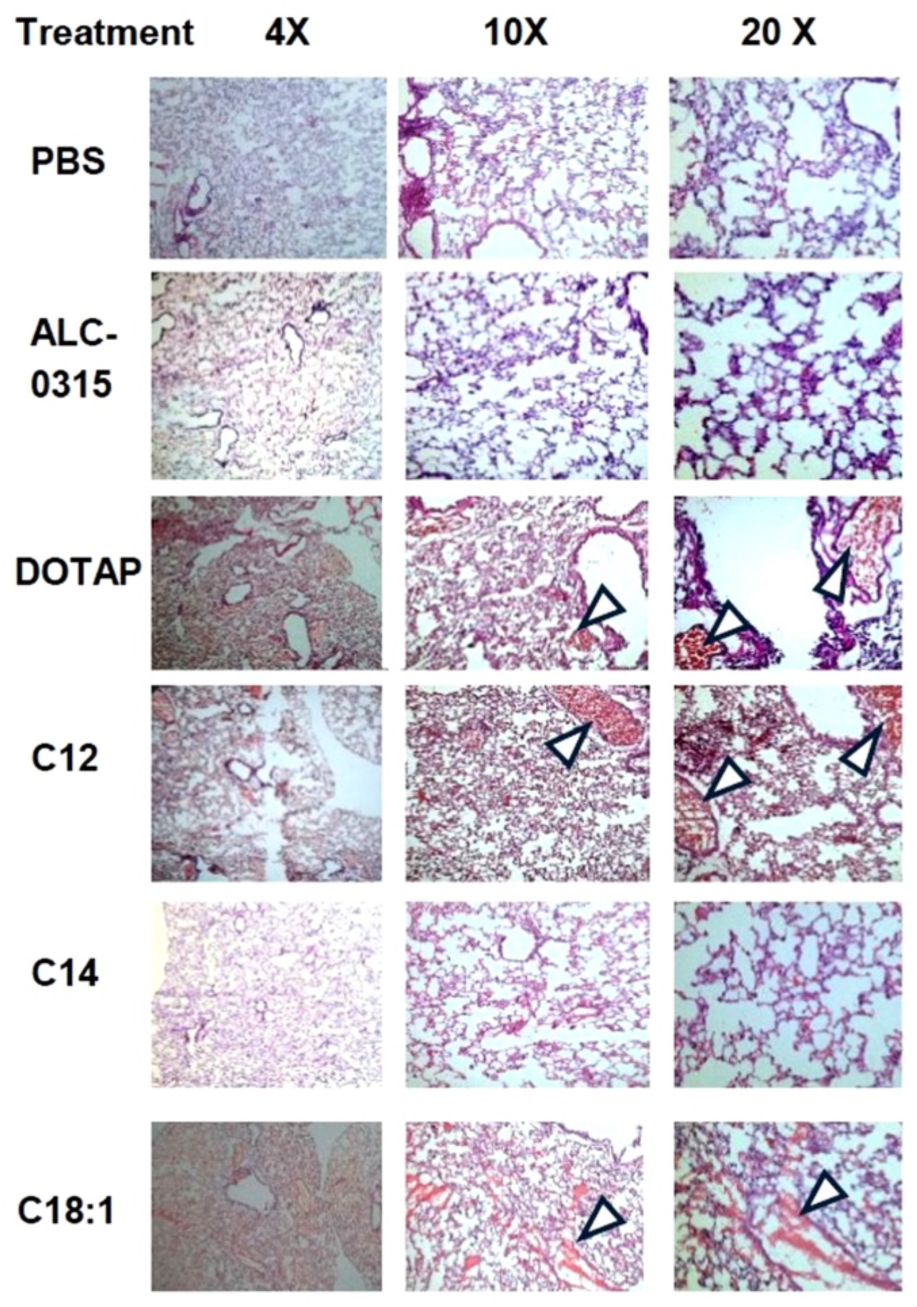
Hematoxylin-eosin (H&E) staining of representative lung sections from mice receiving FLuc-mRNA-LNPs containing either C12, C14, C18:1 TZ lipids or controls. Lung tissues were perfsed and harvested 4 hr post-treatment. Magnification is indicated from left to right as 4x, 10x, and 20x. Histology indicate blood clots in the lungs of mice treated with C12, C18:1 and DOTAP-mRNA LNPs, while mice receiving C14 or ALC-0315 formulations demonstrate similar histology to PBS-treated control mice.

Thrombin-antithrombin (TAT) levels were also measured in the plasma of the TZ lipid-treated and control mice. TAT is a stable complex formed after thrombin activation, acting as a coagulation marker. Notably, the overall TAT concentration in mice treated with the C14 TZ lipid (2.51 ± 0.43 ng/mL) was similar to those treated with ALC-0315 formulations (2.78 ± 0.30 ng/mL) and significantly lower than those treated with the C18:1 TZ lipid (4.06 ± 0.89 ng/mL, \**p*<0.05) and the DOTAP-treated mice (4.06 ± 1.26 ng/mL, \**p*<0.05) (**Figure S3**). Additionally, the TAT concentration is significantly elevated in mice treated with C12 (3.58 ± 0.66 ng/mL, \*\**p*<0.01), C18:1(\*\*\**p*<0.001), and DOTAP (\*\*\**p*<0.001) when compared to control mice treated with PBS (1.31 ± 0.92 ng/mL).

### Calibrated Automated Thrombography (CAT) Assay in both ex vivo and in vivo plasma

Based on the in vivo tissue histology data, the blood clotting properties of the TZ-LNPs were further examined in human blood using the Calibrated Automated Thrombography (CAT) assay, in which thrombin generation is quantified, and the clotting kinetics are determined from the concentration over time. The thrombin generation curve demonstrates the overall time course for thrombin formation in the experimental setting (**Figure 4A**). When comparing the positive and the negative controls, the thrombin generation lag time (indicated by the initiation of the curve above background) starts at approximately 2 min and reaches a maximum at approximately 5 min, while the negative control lag time starts at 6 min and reaches a maximum by 10 min.

**Figure 4:**
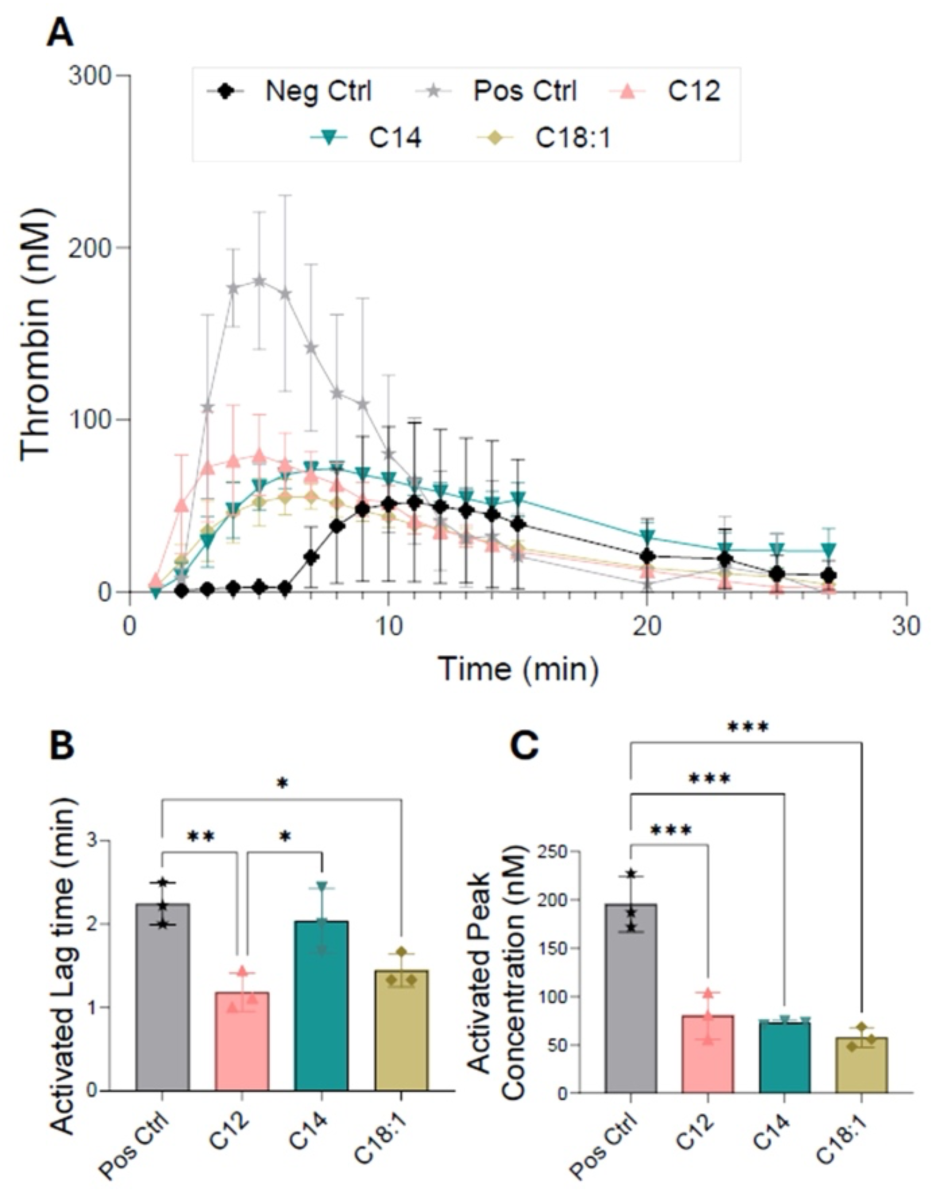
Thrombin generation by thrombin generation assay in pooled human plasma samples incubated with FLuc mRNA-LNPs containing C12, C14, C18:1 TZ lipids. (A) Ex vivo thrombin generation curve for C12, C14, C18:1 TZ lipids as compared to positive and negative controls. (B) Activated lag time quantified as the time required for thrombin formulation to reach detectable levels, (C) activated peak thrombin concentration (nM). Statistical significance is indicated as follows: (*) *p*<0.05, (**) *p*<0.01, (***) *p*<0.001, (****) *p*<0.0001.

When assessing the effects of each LNP formulation on thrombin generation in the activated state in the presence of coagulation factor, the C12 formulation exhibits a significantly shorter onset of thrombin generation (1.2 ± 0.2 min) as compared to both the positive control (2.2 ± 0.3 min, \*\**p* < 0.01) and the C14 TZ lipid (2.0 ± 0.4 min, \**p* < 0.05) (**Figure 4B**). Notably, there are no differences in the unactivated lag time of the formulations (without coagulation factor) (**Figure S4A**). Collectively, these data indicate that C12 and C18:1 TZ lipids, but not the C14 lipid, promote coagulation reactions, accelerating thrombin generation and increasing the potential for clot formation, in alignment with the observed pulmonary thromboses in mice. However, the peak thrombin concentration for each TZ lipid is significantly reduced (<100 nM) as compared to the positive control (195 ± 28 nM), with a similar decrease when coagulation factor is present (activated) (**Figure 4C**) or absent (unactivated) (**Figure S4B**). Although peak thrombin concentration is significantly reduced in the presence of TZ-LNPs, there are no significant differences in total thrombin production (ETP) (**Figure S4C-S4D**). In addition, no statistical differences were observed in the time to reach peak concentration for any of the test lipids (**Figure S4E-S4F**).

## 4. Discussion

In this study, three triazine lipids containing the same head groups with different lipid tail lengths (C12, C14, and C18:1) were synthesized and formulated in LNPs with mRNA by microfluidic mixing, and their activity and safety were assessed in vitro and in vivo. Notably, all three lipids achieve robust protein expression, but have different coagulation risk based on mouse lung pathology and thrombin formation in human plasma. Intriguingly, the C14 saturated lipid tail exhibits minimal thrombotic activity as compared to the other TZ lipids, suggesting that minor structural differences in the lipid architectures can have significant biological implications related to both activity and safety of investigational lipids and formulations.

The mechanism driving differences in protein expression and thrombotic risk are unclear, but the structural differences and resultant physical-chemical characteristics provide insight into potential mechanisms for the observed activity. Formulation attributes such as size, PDI, and surface charge are known to influence cellular uptake and prolonged circulation half-life after systemic administration.^44,45^ However, in this case, the lipid tail lengths do not alter size, PDI, or zeta potential as all TZ formulations exhibit comparable characteristics, suggesting that these attributes have little impact on the observed activity differences between formulations. Additionally, all formulations achieve similar mRNA encapsulation efficiency of over 90%, indicating minimal impact of tail length on mRNA entrapment.

When assessing the formulations for in vitro activity and toxicity, some differences begin to emerge. While in vitro protein expression between the TZ formulations is similar, C12 exhibits increased cytotoxicity as compared to the other TZ formulations, which are similar to the ALC-0315 formulations. The increased cytotoxicity may be due to shorter lipid tail length, which has been reported to be due to loose intermolecular hydrophobic interactions in the cell membrane resulting in disordered packing, and increased cell permeability.^46,47^

Both toxicity and efficacy may also be explained by the p*K*_a_ of the LNP formulations, which influences mRNA protection in the bloodstream, endosomal escape and release of the nucleic acid payload for translation and protein expression.^44^ The optimal LNP p*K*_a_ facilitates protonation in acidic environments of the endo-lysosome for membrane disruption and minimal surface charge in blood to minimize cytotoxicity.^48–50^ The optimal range reported for ionizable lipids that enhance protein expression following intravenous administration is p*K*_a_ = 6.2-6.6.^51^ Of the three TZ lipids, LNPs prepared with the C14 lipid have a p*K*_a_ just above the optimal range with a narrow confidence interval, while C18:1 TZ LNPs exhibit an elevated p*K*_a_ resulting in a cationic formulation across physiologic pH ranges. Given that all three TZ lipids have the same head groups, the differences between p*K*_a_ for each must be driven by the differences in tail length and saturation, which has been reported previously in the presence^52,53^ and absence of nucleic acids.^32^

The relative surface charge and ionizability of the LNP formulations across the physiologic pH range is also of importance for interaction with serum proteins resulting in a protein corona that influences the biodistribution of the formulations. Electrostatic protein interactions with LNPs are influenced partially by the p*K*_a_ of the formulation as well as the ability to interact and remain engaged with the outer leaflet of the LNP membrane.^54–56^ Notably, the C18:1 TZ LNP exhibits a nearly permanent charge across physiologic pH, while the C14 TZ LNP is approaching the ideal p*K*_a_ for an LNP that remains uncharged in circulation and protonated upon acidification in the endo-lysosome. Lung-specific targeting has been reported for LNPs prepared from cationic lipids following systemic administration,^13,21,57,31^ but the mechanisms driving these effects are not fully understood. Correlations between specific proteins in the corona of distinct LNPs and their in vivo biodistribution and activity have been reported, but the engagement between these proteins and different LNPs has not been established.^54,55,58,59^ Nonetheless, LNPs have been shown to accumulate fibrinogen, vitronectin and alpha-1-antitrypsin in their protein coronas, leading to pulmonary-specific delivery.^13,26,29,60^ Importantly, LNPs containing DOTAP, a cationic lipid, have been shown to bind fibrinogen, alter its secondary structure and cause aggregation and activation of platelets.^31^ Therefore, the differences in biodistribution and thromboses between the TZ LNPs may be driven by the minor architectural difference in lipid tail length that modulates p*K*_a_, alters the protein corona and subsequent clotting activity.

The in vivo results of thrombin generation also appear to be magnified as compared to the ex vivo human clotting assay. Although the effects appear to be consistent in mice and ex vivo human plasma, the difference in effect size may be due to the activation of the entire physiological system in mice in vivo, including activation of platelets, which further amplify the clotting cascade. Comparatively, mouse blood coagulates 3-5 times faster in vivo due to the increased number of platelets, and exhibits faster, more reproducible clotting in ex vivo plasma as compared to human blood.^61,62^ The interaction between LNPs and platelets is outside the scope of this work, but considered as a viable target for future experimentation to assess the structure-activity relationships of lipids in human and mouse platelet cell culture. Therefore, it remains unclear if LNPs would cause clotting in humans if administered in the clinic, but a critical safety factor to be investigated for all formulations administered through the intravenous route.

## 5. Conclusion

In summary, this study demonstrates both extrahepatic tissue delivery and risk of pulmonary thrombosis based on minor structural differences in TZ lipid tails. While physicochemical and biophysical interactions are likely to be the cause for the observed findings, continued efforts to dissect the mechanisms of clotting through investigation into the protein corona, platelet activation, and expanded structure-activity relationships are warranted and the focus of future work. Nonetheless, these findings emphasize the effects of minor chemical modifications on thrombotic risk and the necessity to include screening strategies that include thrombin generation when developing new lipid architectures for nucleic acid delivery.

## Acknowledgement

The authors would like to thank Prof. Caroline Geisler and Michaela Murphy from the UK College of Pharmacy for their assistance with cryo-sectioning of the lung tissues. In addition, Melissa Jauquet (formerly at UK College of Medicine) from Texas A&M University helped with the CAT assay.

