## Supplemental Information for "Tail length of triazine-based lipids influences blood clotting risk in vitro and in vivo"

**TZ lipid synthesis steps**

**Note**: Peak-broadening in the $^{1}H$ and $^{13}C\{^{1}H\}$ NMR spectra of di- and tri-substituted 1,3,5-triazines was observed, which is consistent with restricted rotation around the C–N bonds connecting the triazine carbon atoms to the external amines.

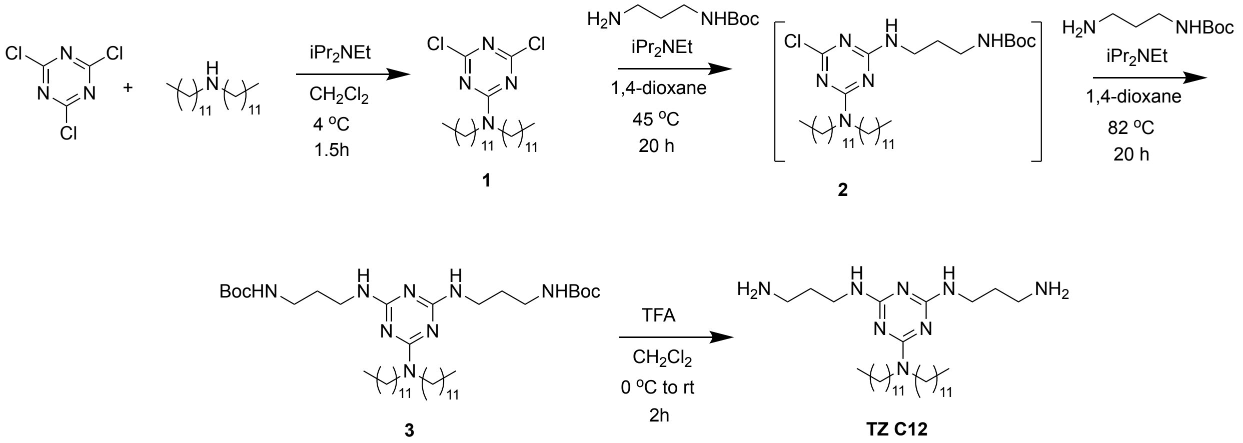

**Scheme S1.** Synthetic scheme for Lipid TZ C12.

**Compound 1**.[2] Cyanuric chloride (1.88 g, 10.2 mmol) was added to a stirred solution of didodecylamine (3.00 g, 8.48 mmol) in 50 mL of dichloromethane at 4 °C under air. Diisopropylethylamine (7.4 mL, 5.5 g, 42.4 mmol) was added slowly, and the reaction stirred at 4 °C. After 1.5 h, the reaction solution warmed to rt and was washed with 30 mL of 1M aqueous acetic acid and 30 mL of saturated aqueous sodium chloride. The organic layer was dried over anhydrous sodium sulfate, filtered, and evaporated under reduced pressure. The resulting tan solid was suspended in 10 mL of boiling methanol, and a minimal amount of chloroform was added to dissolve it. Upon cooling to 4 °C, a white solid formed. It was collected by vacuum filtration and washed with 3 x 5 mL of 4 °C methanol to afford 3.83 g (90%) of **1** as a white solid. Spectroscopic data was consistent with reported values.[2] $^{1}H$ NMR (400 MHz, $CDCl_3$, ppm): $\delta$ 3.53 (t, $J$ = 7.6 Hz, 4H), 1.59 (m, 4H), 1.26–1.31 (m, 36H), 0.88 (t, $J$ = 6.8 Hz, 6H).$^{13}C\{^{1}H\}$NMR (100 MHz, $CDCl_3$, ppm): $\delta$ 169.77, 164.29, 47.85, 31.92, 29.65, 29.58, 29.50, 29.35, 29.25, 27.16, 26.69, 22.69, 14.12. HRMS MW calculated for $C_{27}H_{50}Cl_2N_4$ $(M+H)^+$: = 501.3485; found: 501.3542.

**Compound 2**. A solution of *N*-(3-aminopropyl)carbamic acid *t*-butyl ester (0.54 g, 3.1 mmol) in 4 mL of 1,4-dioxane was added to a solution of **1** (1.3 g, 2.6 mmol) in 18 mL of 1,4-dioxane at rt. Diisopropylethylamine (2.3 mL, 1.68 g, 13.0 mmol) was added to the reaction solution, which was heated to 45 °C with stirring. After 20 h, the volatiles were removed under reduced pressure. The remaining residue was dissolved in 40 mL of dichloromethane and was washed with 2 x 40 mL of saturated aqueous sodium chloride. The organic layer was dried over anhydrous sodium sulfate, filtered, and evaporated. The resulting solid was purified by recrystallization from 6 mL of methanol to afford 1.53 g (82%) of **2** as a white solid. $^{1}$H NMR (400 MHz, $CDCl_3$, ppm): *δ* 3.45–3.49 (m, 6H), 3.40–3.43 (br q, *J* = 6.0 Hz, 2H), 1.71–1.73 (m, 2H), 1.57 (m, 4H), 1.44 (s, 9H), 1.26 (m, 36H), 0.88 (t, *J* = 6.8 Hz, 6H).$^{13}$C{$^{1}$H}NMR (100 MHz, $CDCl_3$, ppm): *δ* 168.46, 165.45, 164.53, 156.14, 79.22, 47.46, 47.24, 46.94, 37.8, 37.65, 31.92, 29.94, 29.67, 29.64, 29.58, 29.49, 29.40, 29.36, 28.40, 27.85, 27.51, 27.06, 26.80, 22.69, 14.12. HRMS MW calculated for $C_{35}H_{67}ClN_6O_2$ $(M+H)^+$: = 639.5087; found: 639.4431.

**Compound 3**. Diisopropylethylamine (3.7 mL, 2.74 g, 21.2 mmol) was added to a solution of **2** (1.35 g, 2.12 mmol) and *N*-(3-aminopropyl)carbamic acid *t*-butyl ester (1.84 g, 10.6 mmol) in 20 mL of 1,4-dioxane. After stirring at 82 °C for 2 days, the volatiles were removed under reduced pressure. The remaining oily residue was dissolved in 50 mL of dichloromethane and washed with 2 x 30 mL of water and 30 mL of saturated aqueous sodium chloride. The organic layer was dried over anhydrous sodium sulfate, filtered, and evaporated to an oil that was purified by flash chromatography (3% methanol/97% dichloromethane). The resulting white solid was triturated with 4 °C methanol to afford 1.10 g (67%) of **3** as a white solid. $R_f$ = 0.30 in 4% methanol/96% dichloromethane. $^{1}$H NMR (400 MHz, $CDCl_3$, ppm): *δ* 3.43–3.49 (m, 8H), 3.17–3.19 (m, 4H), 1.69 (m, 4H), 1.56 (m, 4H), 1.44 (s, 18H), 1.26–1.28 (m, 36H), 0.86–0.90 (t, *J* = 6.8 Hz, 6H). $^{13}$C{$^{1}$H}NMR (100 MHz, $CDCl_3$, ppm): *δ* 164.90, 161.55, 156.12, 79.05, 39.63, 38.40, 33.31, 31.93, 29.69, 29.68, 29.66, 29.56, 29.37, 28.44, 28.05, 27.11, 22.69, 14.13. HRMS MW calculated for $C_{43}H_{84}N_8O_4$ $(M+H)^+$: = 777.6688; found: 777.6757.

**Lipid TZ C12**.[2] Trifluoroacetic acid (16.3 mL, 24.2 g, 212 mmol) was added to a stirred solution of **3** (1.10 g, 1.42 mmol) in 16 mL of dichloromethane at 4 °C. The solution warmed to rt, and after 2 h, the volatiles were removed under reduced pressure. The remaining oily residue was dissolved in 15 mL of dichloromethane and was washed with 2 x 15 mL of 2M aqueous sodium hydroxide. The combined aqueous layers were extracted with 10 mL of dichloromethane, and the combined organic layers were dried over anhydrous sodium sulfate, filtered, and evaporated to afford 555 mg (68%) of **TZ C12** as a pale-yellow oil. Spectroscopic data was consistent with reported values.[2] $^{1}$H NMR (400 MHz, $CDCl_3$, ppm): *δ* 3.40–3.48 (m, 8H), 2.78 (t, *J* = 6.8 Hz, 4H), 1.68 (quint, *J* = 6.8 Hz, 4H), 1.51–1.60 (m, 4H), 1.26–1.30 (m, 36H), 0.88 (t, *J* = 6.8 Hz, 6H). $^{13}$C{$^{1}$H}NMR (100 MHz, $CDCl_3$, ppm): *δ* 166.16, 165.00, 46.78, 39.60, 38.09, 33.59, 31.93, 29.70, 29.668, 29.66, 29.56, 29.37, 28.06, 27.14, 22.80, 22.69, 14.12. HRMS MW calculated for $C_{33}H_{68}N_8$ $(M+H)^+$: = 577.5640; found: 577.5099.

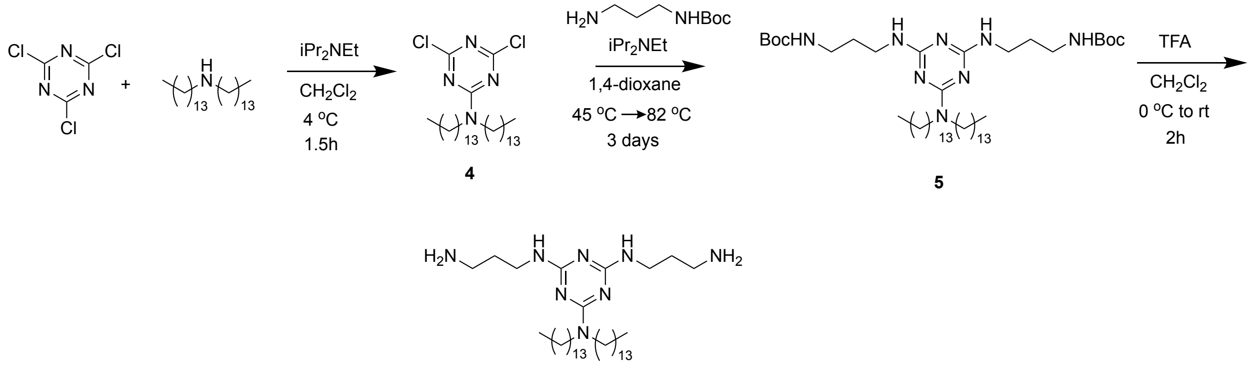


**Scheme S2.** Synthetic scheme for lipid TZ C14.

**Compound 4**. Cyanuric chloride (0.54 g, 2.9 mmol) was added to a stirred solution of ditetradecylamine (1.00 g, 2.44 mmol) in 25 mL of dichloromethane at 4 °C under air. Diisopropylethylamine (2.1 mL, 1.58 g, 12.2 mmol) was added slowly, resulting in a slurry that stirred at 4 °C. After 1.5 h, the reaction solution warmed to rt and was diluted with 40 mL of dichloromethane. It was washed with 35 mL of 1M aqueous acetic acid and 35 mL of saturated aqueous sodium chloride. The organic layer was dried over anhydrous sodium sulfate, filtered, and evaporated under reduced pressure. The resulting pale-yellow solid was triturated with 5 mL of cold methanol and was collected by vacuum filtration and washed with 3 x 4 mL of 4 °C methanol to afford 1.19 g (88%) of **4** as a white solid. $^{1}H$ NMR (400 MHz, $CDCl_3$, ppm): $\delta$ 3.53 (t, $J$ = 7.6 Hz, 4H), 1.59–1.61 (m, 4H), 1.26–1.30 (m, 44H), 0.88 (t, $J$ = 6.8 Hz, 6H). $^{13}C\{^{1}H\}$NMR (100 MHz, $CDCl_3$, ppm): $\delta$ 169.79, 164.32, 47.87, 31.94, 29.70, 29.68, 29.66, 29.65, 29.59, 29.50, 29.37, 29.25, 27.17, 26.70, 22.70, 14.12. HRMS MW calculated for $C_{31}H_{58}Cl_2N_4$ $(M+H)^+$: = 557.4111; found: 557.4177.

**Compound 5**. A solution of *N*-(3-aminopropyl)carbamic acid *t*-butyl ester (375 mg, 2.15 mmol) in 3 mL of 1,4-dioxane was added to a mixture of **4** (1.00 g, 1.79 mmol) in 10 mL of 1,4-dioxane at rt. Diisopropylethylamine (1.6 mL, 1.16 g, 8.97 mmol) was added to the reaction mixture, which was heated to 45 °C with stirring. After 20 h, the reaction solution was cooled to rt, and more *N*-(3-aminopropyl)carbamic acid *t*-butyl ester (1.56 g, 8.97 mmol) and diisopropylethylamine (1.9 mL, 1.39 g, 10.8 mmol) were added. After stirring at 82 °C for two days, the volatiles were removed under reduced pressure. The remaining oily residue was dissolved in 35 mL of dichloromethane and washed with 2 x 35 mL of water and 35 mL of saturated aqueous sodium chloride. The organic layer was dried over anhydrous sodium sulfate, filtered, and evaporated to an oil that was purified by flash chromatography (3% methanol/97% dichloromethane). The resulting viscous yellow oil was triturated with 4 °C methanol to afford 1.06 g (71%) of **5** as a white solid. $R_f$ = 0.20 in 3% methanol/97% dichloromethane. $^{1}H$ NMR (400 MHz, $CDCl_3$, ppm): $\delta$

3.40–3.46 (m, 8H), 3.17–3.18 (m, 4H), 1.67–1.71 (m, 4H), 1.52–1.58 (m, 4H), 1.44 (s, 18H), 1.26–1.28 (m, 44H), 0.88 (t, $J$ = 6.8 Hz, 6H). $^{13}C\{^1H\}$NMR (100 MHz, $CDCl_3$, ppm): $\delta$ 164.87, 156.10, 78.88, 46.94, 37.36, 31.93, 30.42, 29.70, 29.66, 29.56, 29.37, 28.47, 28.05, 27.12, 26.75, 22.69, 14.12. HRMS MW calculated for $C_{47}H_{92}N_8O_4$ $(M+H)^+$: = 833.7314; found: 833.6480.

**Lipid TZ C14**. Trifluoroacetic acid (4.9 mL, 7.23 g, 63.4 mmol) was added to a stirred solution of **5** (352 mg, 0.42 mmol) in 4.9 mL of dichloromethane at 4 °C. The solution warmed to rt, and after 2 h, the volatiles were removed under reduced pressure. The remaining oily residue was dissolved in 5 mL of dichloromethane and was washed with 2 x 5 mL of 2M aqueous sodium hydroxide. The combined aqueous layers were extracted with 3 mL of dichloromethane, and the combined organic layers were dried over anhydrous sodium sulfate, filtered, and evaporated to afford 193 mg (72%) of **TZ C14** as a pale-yellow oil. $^1H$ NMR (400 MHz, $CDCl_3$, ppm): $\delta$ 3.40–3.48 (m, 8H), 2.79 (t, $J$ = 6.8 Hz, 4H), 1.65–1.72 (m, 4H), 1.51–1.59 (m, 4H), 1.25–1.30 (m, 44H), 0.88 (t, $J$ = 6.8 Hz, 6H). $^{13}C\{^1H\}$NMR (600 MHz, $CDCl_3$, ppm): $\delta$ 165.87, 164.91, 164.64, 46.82, 39.31, 38.00, 32.90, 31.94, 29.72, 29.70, 29.57, 29.38, 28.07, 27.15, 22.81, 22.70, 22.58, 14.13. HRMS MW calculated for $C_{37}H_{76}N_8$ $(M+H)^+$: = 633.6266; found: 633.5697.

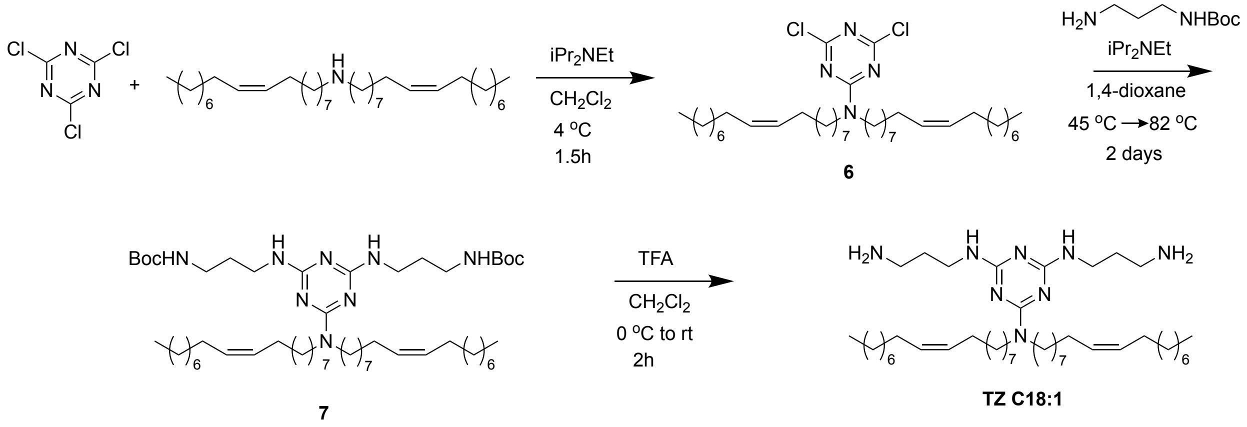


**Scheme S3.** Synthetic scheme for lipid TZ C18:1.

**Compound 6**. Cyanuric chloride (871 mg, 4.72 mmol) was added to a stirred solution of dioleylamine[1] (2.04 g, 3.93 mmol) in 25 mL of dichloromethane at 4 °C under air. Diisopropylethylamine (3.4 mL, 2.54 g, 19.7 mmol) was added slowly, and the reaction stirred at 4 °C. After 1.5 h, the reaction solution warmed to rt and was washed with 25 mL of 1M aqueous acetic acid and 25 mL of saturated aqueous sodium chloride. The organic layer was dried over anhydrous sodium sulfate, filtered, and evaporated under reduced pressure. The resulting yellow oil was purified by flash chromatography (2%

ethyl acetate/98% cyclohexane) to afford 2.33 g (89%) of **6** as a pale-yellow oil. $R_f$ = 0.33 in 2% ethyl acetate/98% cyclohexane. $^1$H NMR (400 MHz, $CDCl_3$, ppm): *δ* 5.34–5.38 (m, 4H), 3.53 (t, *J* = 7.6 Hz, 4H), 1.99–2.02 (m, 8H), 1.59–1.61 (m, 4H), 1.27–1.31 (m, 44H), 0.88 (t, *J* = 6.8 Hz, 6H). $^{13}$C{$^1$H}NMR (100 MHz, $CDCl_3$, ppm): *δ* 169.80, 164.33, 129.99, 129.74, 47.85, 31.91, 29.77, 29.72, 29.53, 29.39, 29.32, 29.21, 27.23, 27.18, 26.69, 22.68, 14.10. HRMS MW calculated for $C_{39}H_{70}Cl_2N_4$ $(M+H)^+$: = 665.5050; found: 665.4505.

**Compound 7**. A solution of *N*-(3-aminopropyl)carbamic acid *t*-butyl ester (314 mg, 1.80 mmol) in 3 mL of 1,4-dioxane was added to a mixture of **6** (1.00 g, 1.50 mmol) in 9 mL of 1,4-dioxane at rt. Diisopropylethylamine (1.3 mL, 0.97 g, 7.51 mmol) was added to the reaction mixture, which was heated to 45 °C with stirring. After 20 h, the reaction solution was cooled to rt, and more *N*-(3-aminopropyl)carbamic acid *t*-butyl ester (1.31 g, 7.51 mmol) and diisopropylethylamine (1.6 mL, 1.16 g, 9.0 mmol) were added. After stirring at 82 °C for two days, the volatiles were removed under reduced pressure. The remaining oily residue was dissolved in 30 mL of dichloromethane and washed with 2 x 25 mL of saturated aqueous ammonium chloride, 25 mL of water, and 25 mL of saturated aqueous sodium chloride. The organic layer was dried over anhydrous sodium sulfate, filtered, and evaporated to a brown oil that was purified by flash chromatography (3% methanol/97% dichloromethane) to afford 1.12 g (79%) of **7** as a viscous, pale-yellow oil. $R_f$ = 0.21 in 3% methanol/97% dichloromethane. $^1$H NMR (400 MHz, $CDCl_3$, ppm): *δ* 5.30–5.38 (m, 4H), 4.99, (3.40–3.43 (m, 8H), 3.16–3.18 (m, 4H), 1.97–2.02 (m 8H), 1.67–1.70 (m, 4H), 1.52–1.58 (m, 4H), 1.44 (s, 18H), 1.27–1.29 (m, 44H), 0.88 (t, *J* = 7.2 Hz, 6H). $^{13}$C{$^1$H}NMR (100 MHz, $CDCl_3$, ppm): *δ* 164.81, 161.70, 156.10, 129.93, 129.81, 78.90, 47.01, 37.34, 32.61, 31.90, 30.37, 29.78, 29.77, 29.71, 29.66, 29.59, 29.53, 29.32, 28.47, 28.41, 28.37, 28.05, 27.21, 27.12, 22.68, 14.12. HRMS MW calculated for $C_{55}H_{104}N_8O_4$ $(M+H)^+$: = 941.8253; found: 941.7418.

**Lipid TZ C18:1**. Trifluoroacetic acid (9.1 mL, 13.5 g, 118.7 mmol) was added to a stirred solution of **7** (745 mg, 0.79 mmol) in 9.1 mL of dichloromethane at 4 °C. The solution warmed to rt, and after 2 h, the volatiles were removed under reduced pressure. The remaining oily residue was dissolved in 15 mL of ethyl acetate and was washed with 2 x 15 mL of 2M aqueous sodium hydroxide. The combined aqueous layers were extracted with 10 mL of ethyl acetate, and the combined organic layers were dried over anhydrous sodium sulfate, filtered, and evaporated to afford 328 mg (56%) of **TZ C18:1** as a yellow oil. $^1$H NMR (400 MHz, $CDCl_3$, ppm): *δ* 5.27–5.31 (m, 4H), 4.87, 3.33–3.40 (m, 8H), 2.71 (t, *J* = 6.8 Hz, 4H), 1.89–2.00 (m, 8H), 1.61–1.64 (quint, *J* = 6.4 Hz, 4H), 1.43–1.51 (m, 4H), 1.16–1.34 (m, 44H), 0.80 (t, *J* = 6.8 Hz, 6H). $^{13}$C{$^1$H}NMR (100 MHz, $CDCl_3$, ppm): *δ* 164.97, 163.86, 128.90, 128.78, 45.79, 38.14, 36.89, 36.79, 36.52, 31.58, 30.87, 28.75, 28.74, 28.68, 28.64, 28.58, 28.50, 28.29, 28.16, 27.05, 26.19, 26.12, 25.98, 24.71, 24.64, 21.65, 13.08. HRMS peak resolution remained insufficient despite multiple optimization attempts using varying analyte concentrations and mobile-phase gradient conditions.

**Table S1:** Composition of lipid nanoparticles tested

| **Samples** | **Composition** | **Mol ratio** |
|---|---|---|
| C12 | Lipid TZ C12 : DOPE : DSPE-PEG2000 : Chol | 50 : 10 : 1.5 : 38.5 |
| C14 | Lipid TZ C14 : DOPE : DSPE-PEG2000 : Chol | 50 : 10 : 1.5 : 38.5 |
| C18:1 | Lipid TZ C18:1 : DOPE : DSPE-PEG2000 : Chol | 50 : 10 : 1.5 : 38.5 |
| ALC-0315 | ALC-0315 : DSPC : DMA-PEG2000 : Chol | 46 : 9.4 : 1.7 : 42.9 |
| DOTAP | DOTAP : CKK-E12 : DOPE : DMG-PEG2000 : Chol | 50 : 25 : 5 : 1.5 : 18.5 |

**Table S2:** p$K_a$ of lipid nanoparticles tested by TNS assay

| Lipid with FLuc mRNA | p$K_a$ |
|---|---|
| C12 | 8.85 ± 7.38 |
| C14 | 7.07 ± 0.91 |
| C18:1 | 9.0 ± 1.63 |
| Alc-0315 | 6.41 ± 0.28 |
| DOTAP | Undetectable |

| **Lipids** | **Ex vivo imaging of mice organs** | |
|---|---|---|
| ALC-0315 | 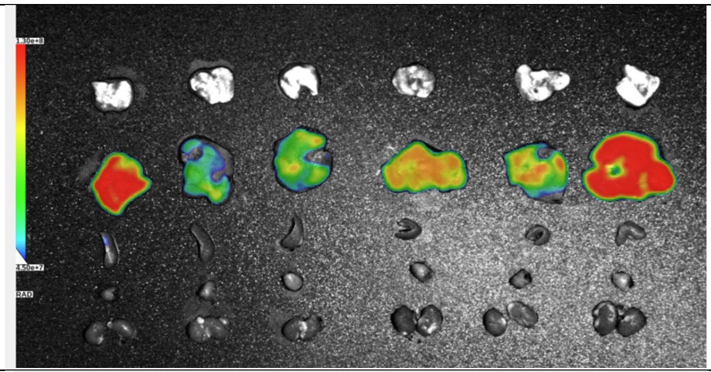 | Lung<br>Liver<br>Spleen<br>Heart<br>Kidney |
| DOTAP | 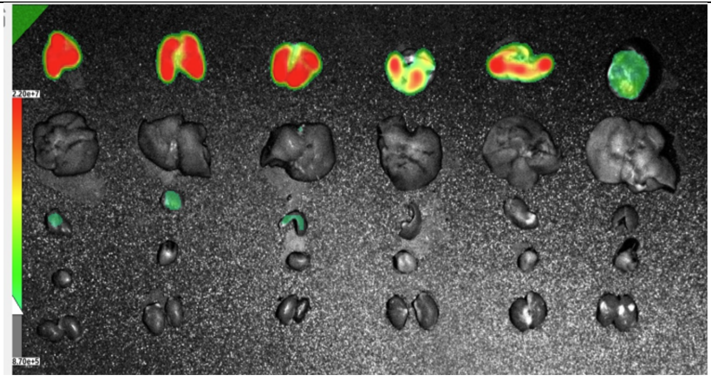 | Lung<br>Liver<br>Spleen<br>Heart<br>Kidney |
| C12 | 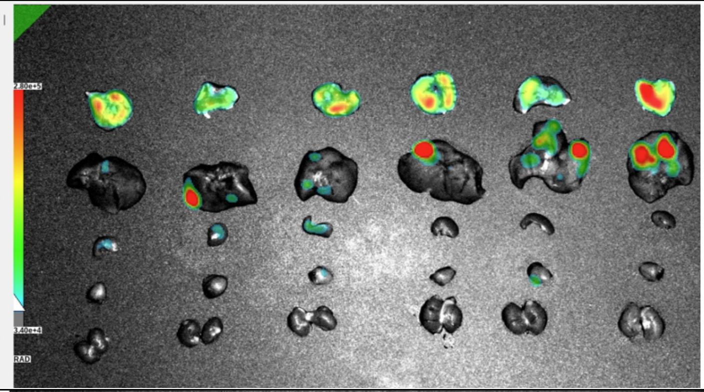 | Lung<br>Liver<br>Spleen<br>Heart<br>Kidney |
| C14 | 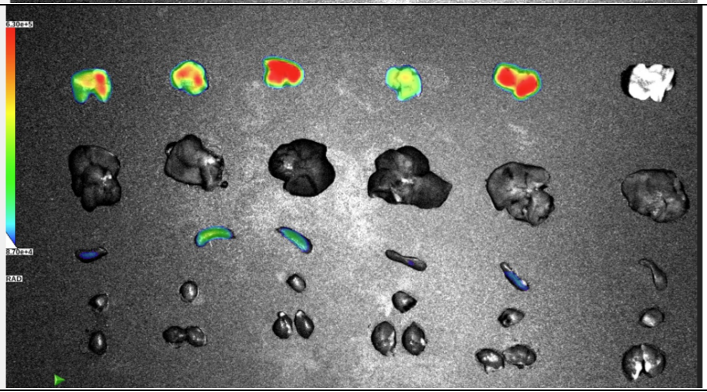 | Lung<br>Liver<br>Spleen<br>Heart<br>Kidney |

| Lipids | Ex vivo imaging of mice organs | |
|---|---|---|
| C18:1 | 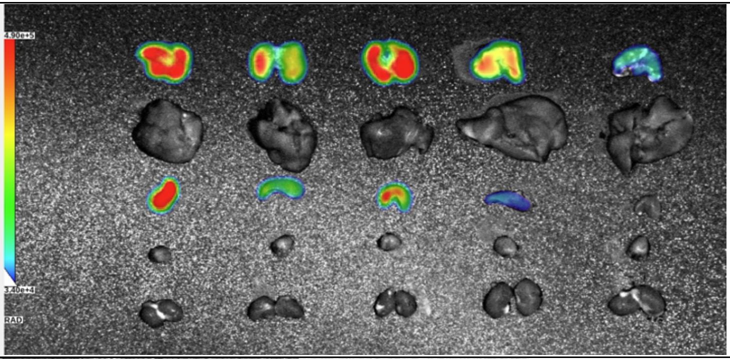 | Lung<br>Liver<br>Spleen<br>Heart<br>Kidney |
| PBS | 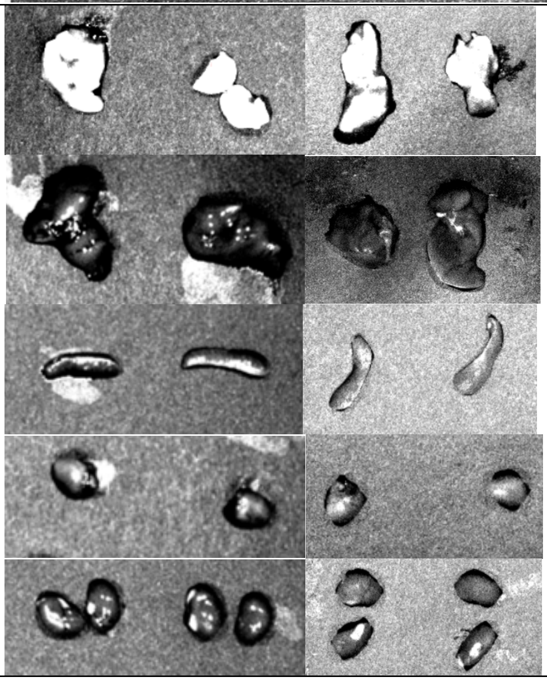 | Lung<br>Liver<br>Spleen<br>Heart<br>Kidney |

**Figure S1**: In vivo biodistribution and FLuc mRNA expression features of LNPs formulated with C12, C14, C18:1 TZ lipids, following intravenous administration. The ex vivo bioluminescence images of major organs (lung, liver, spleen, heart and kidney) (n = 6). PBS was used as control (n=4).

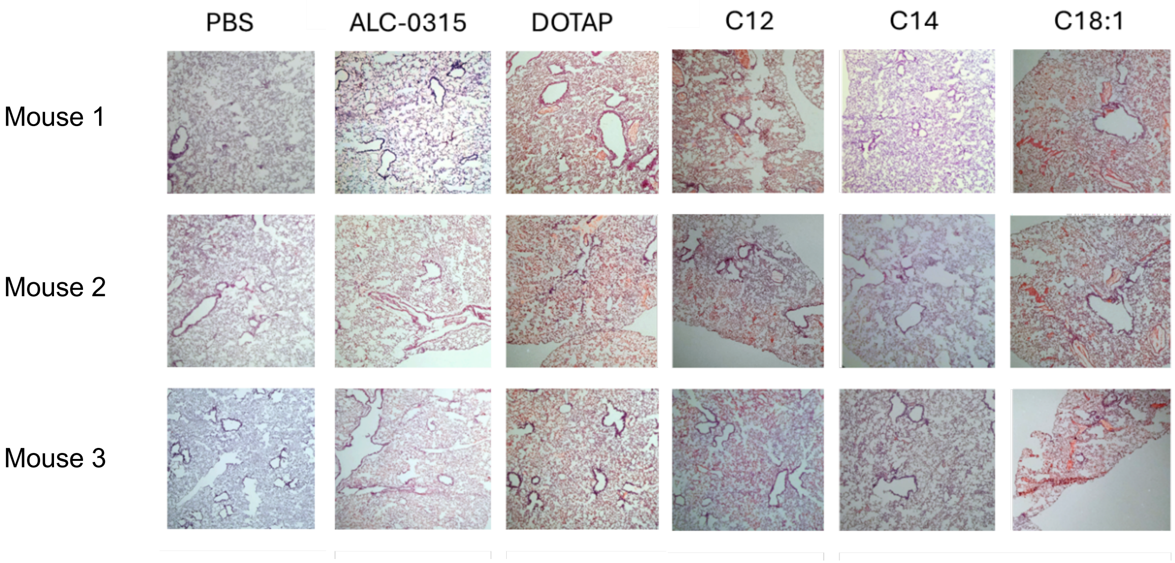


**Figure S2**: Hematoxylin-eosin (H & E) staining of lung sections at 4X magnification from mice receiving different formulations of FLuc mRNA-LNPs containing C12, C14, C18:1 TZ lipids. Lung tissues were perfused and harvested 4 h post-treatment. Histology showed blood clots in the lungs of mice treated with C12, C18:1, and DOTAP-mRNA LNPs. However, mice receiving C14 and ALC-0315 formulations showed a similar morphology to PBS-treated control mice.

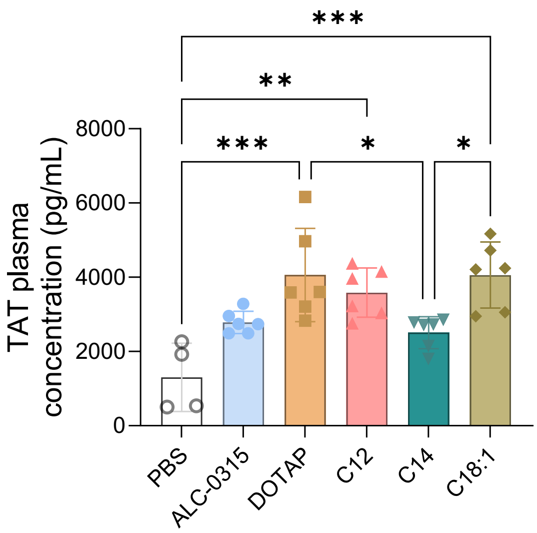


**Figure S3**: Thrombin generation was measured in plasma samples collected from mice treated with 0.25 mg/kg dose of FLuc mRNA-LNPs. Concentration of thrombin-antithrombin complex formed in mice plasma (n=4 to 6) was determined by kit-based ELISA. Data are considered significant (*) where *p*-value <0.05. **p<0.01, ***p<0.001, ****p<0.0001.

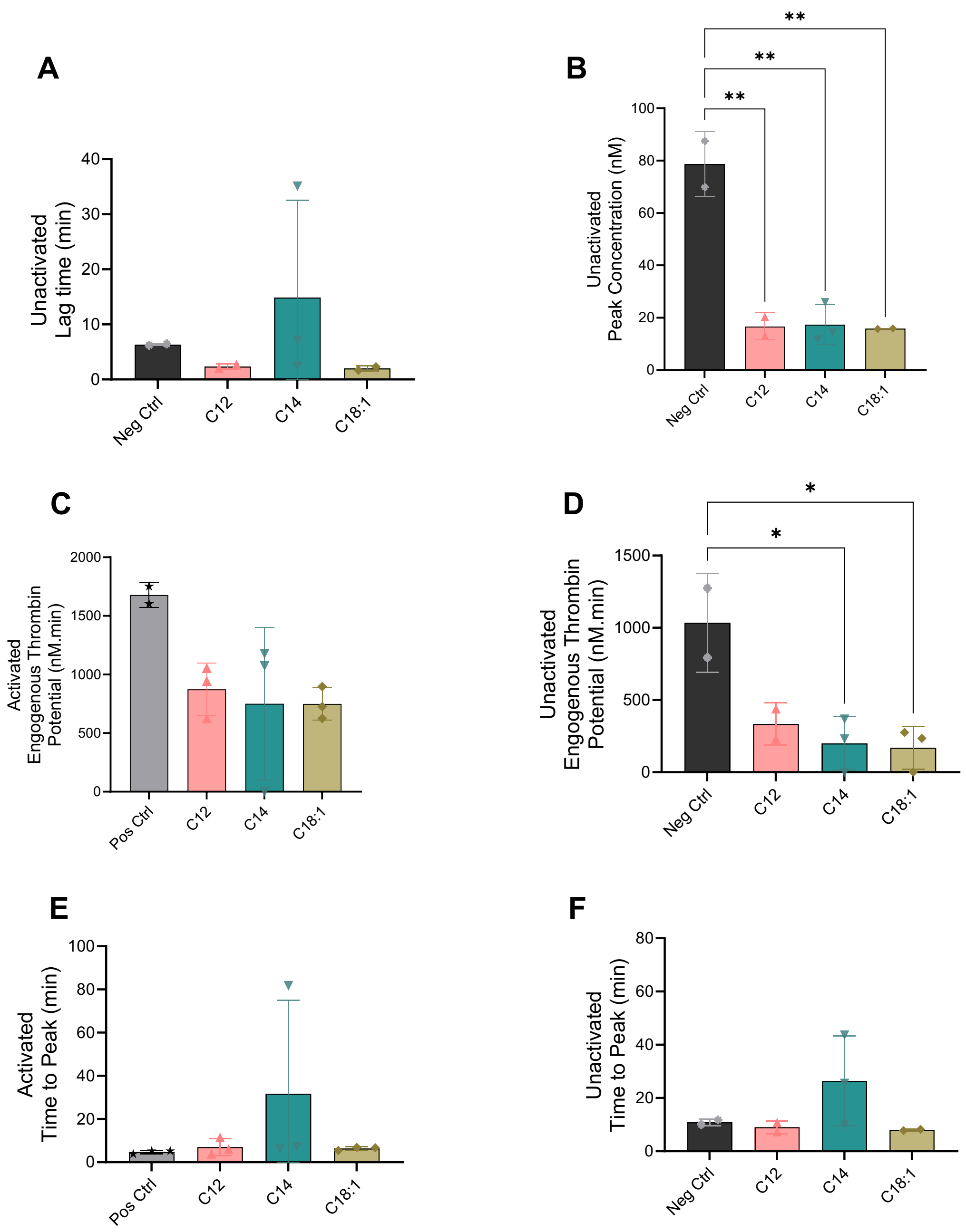


**Figure S4**: CAT assay in pooled human plasma. (A) Unactivated lag time (n=2 to 3), (B) Unactivated peak concentration (n=2 to 3), (C) Activated endogenous thrombin potential (n=2 to 3), (D) Unactivated endogenous thrombin potential (n=2 to 3), (E) Activated time to peak (n =3). (F) Unactivated time to peak (n=2 to 3).

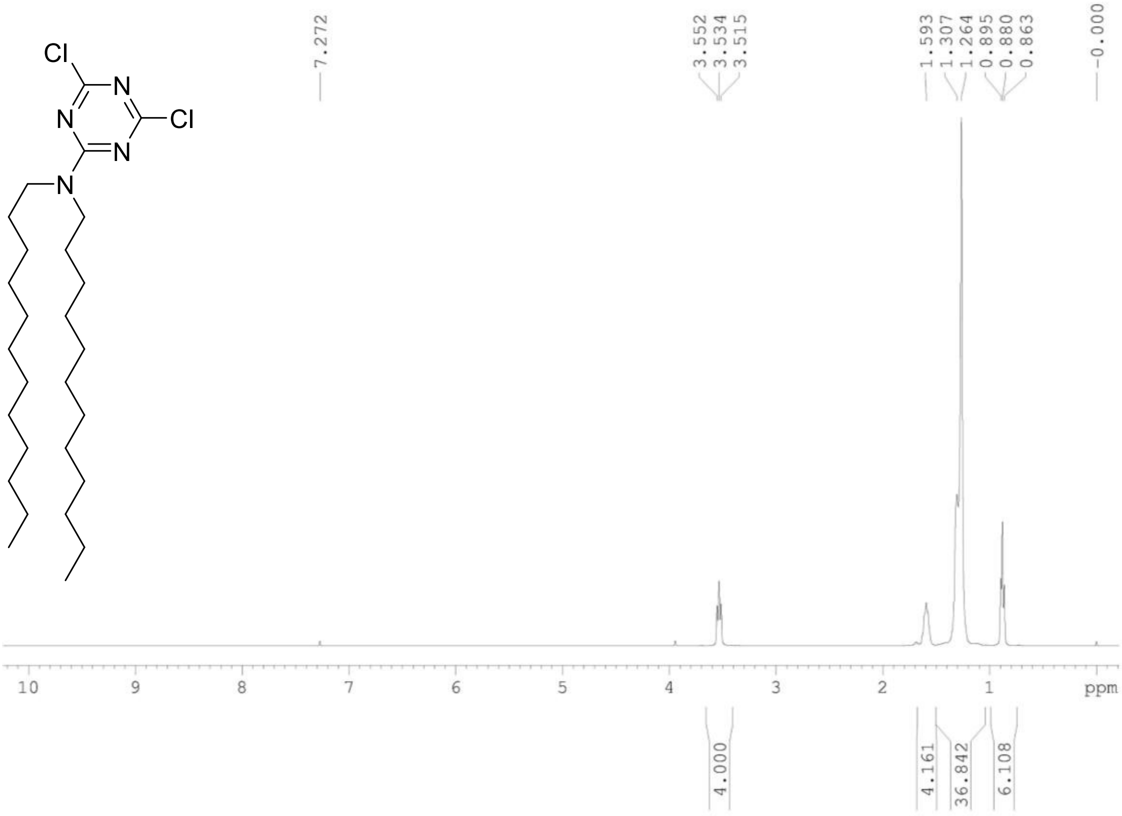


**Figure S5A:** $^{1}$H NMR Spectra of Compound 1

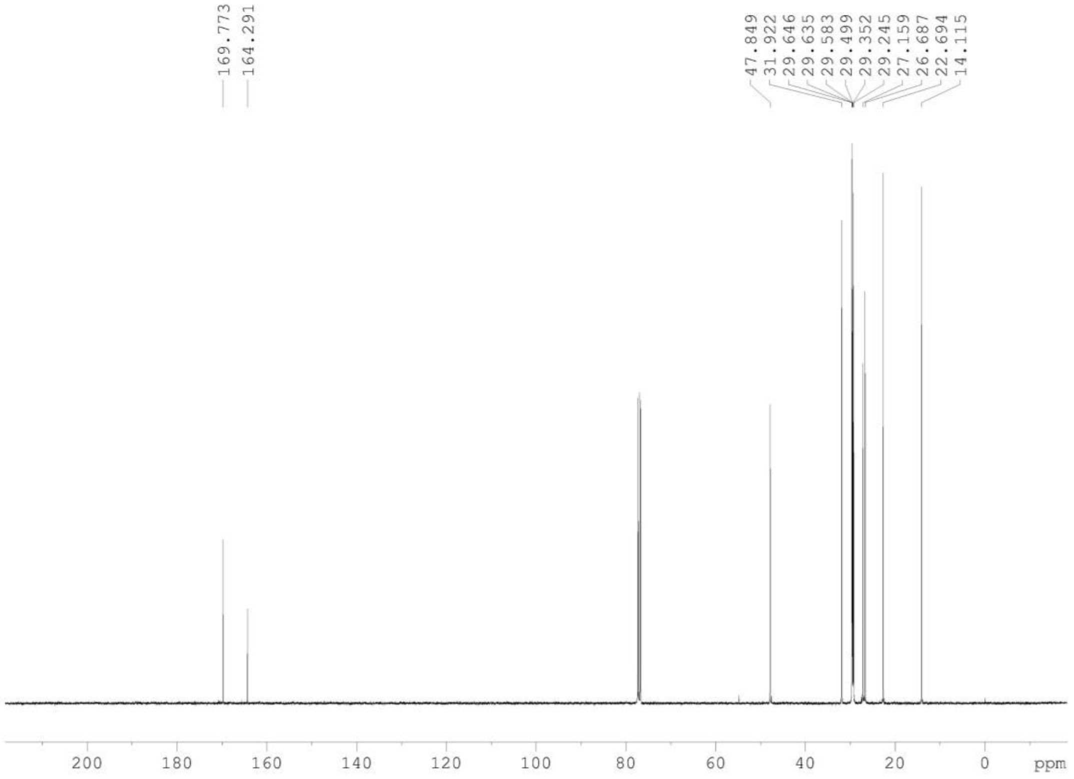


**Figure S5B:** $^{13}$C NMR Spectra of Compound 1

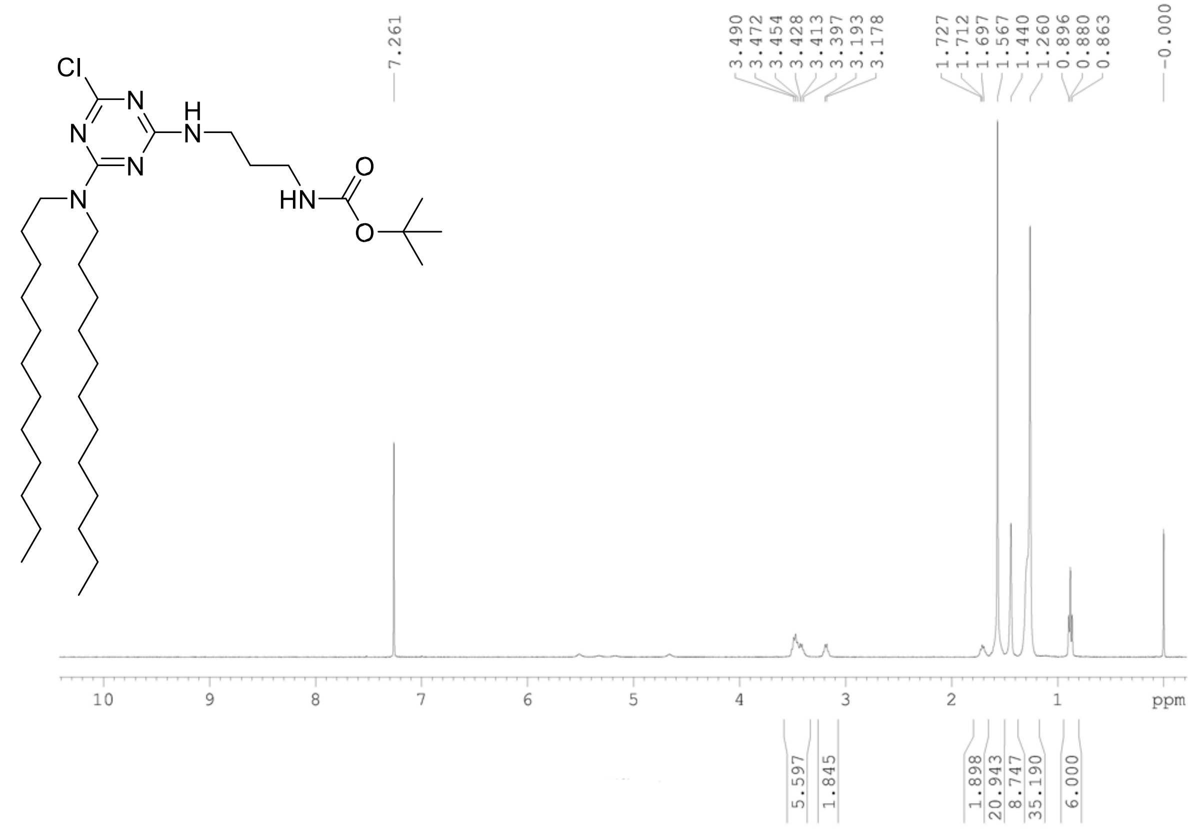


**Figure S6A:** $^{1}$H NMR Spectra of Compound 2

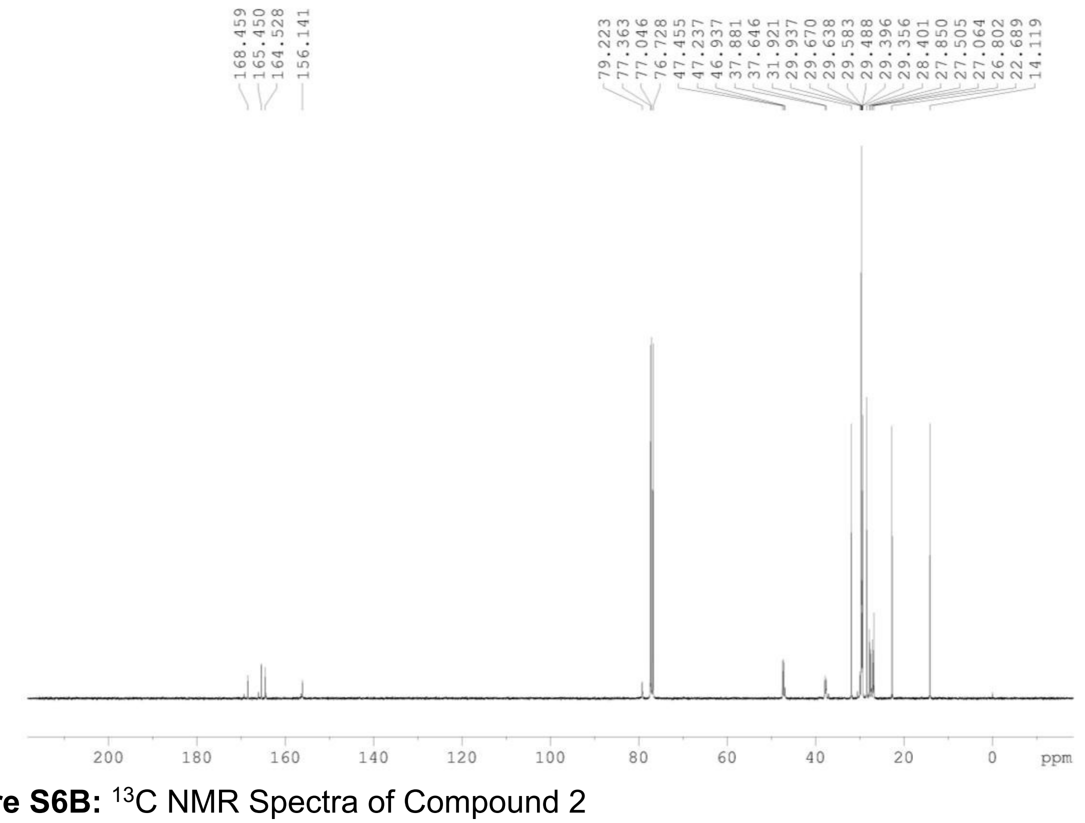


**Figure S6B:** $^{13}$C NMR Spectra of Compound 2

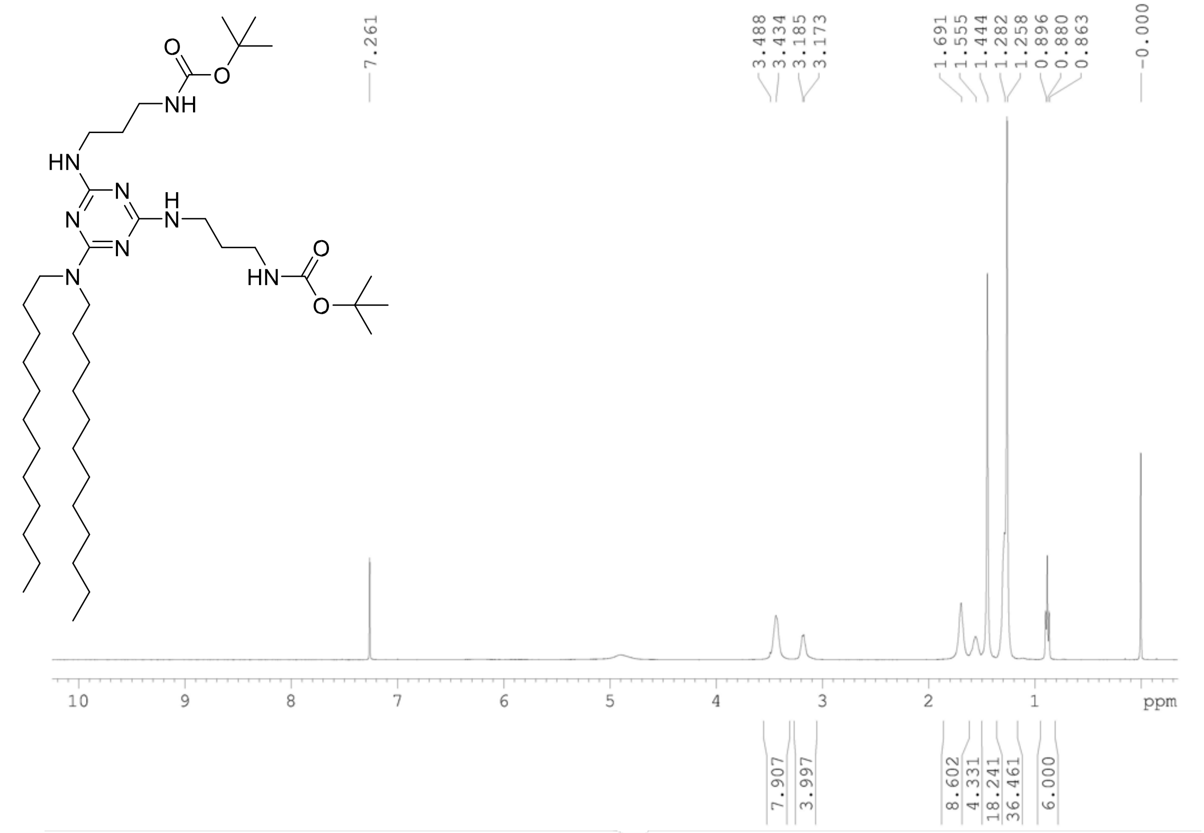


**Fig S7A:** $^{1}$H NMR Spectra of Compound 3

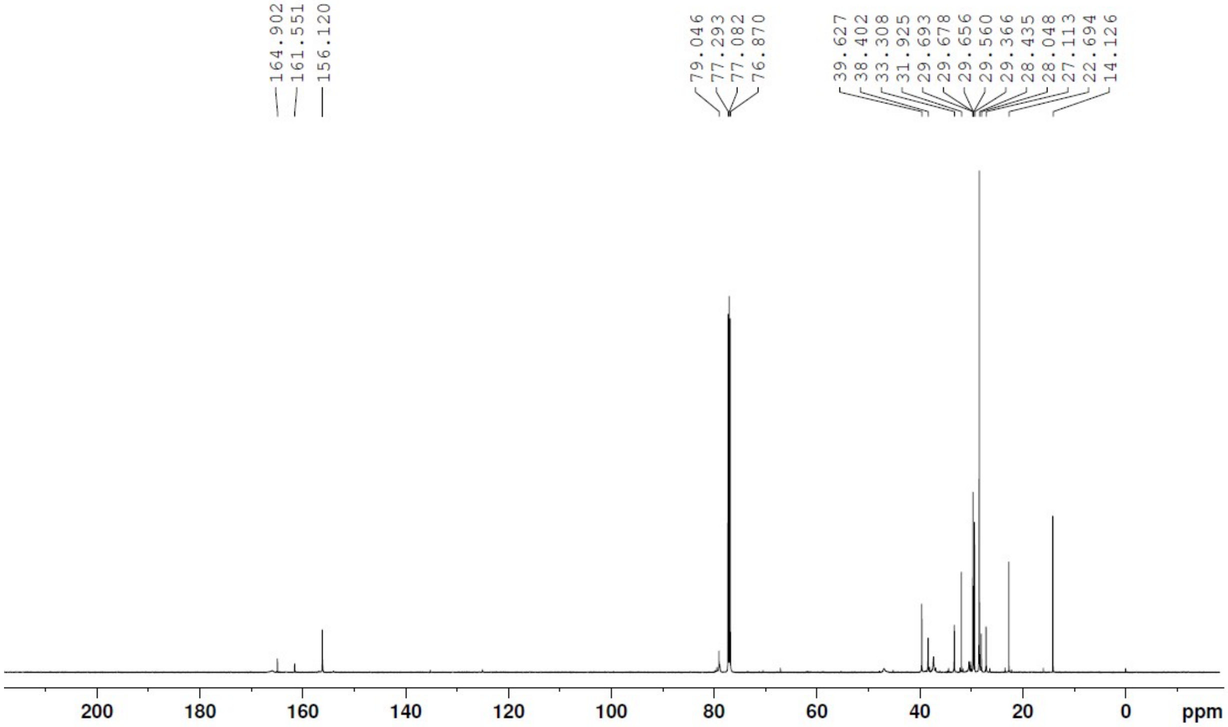


**Fig S7B:** $^{13}$C NMR Spectra of Compound 3

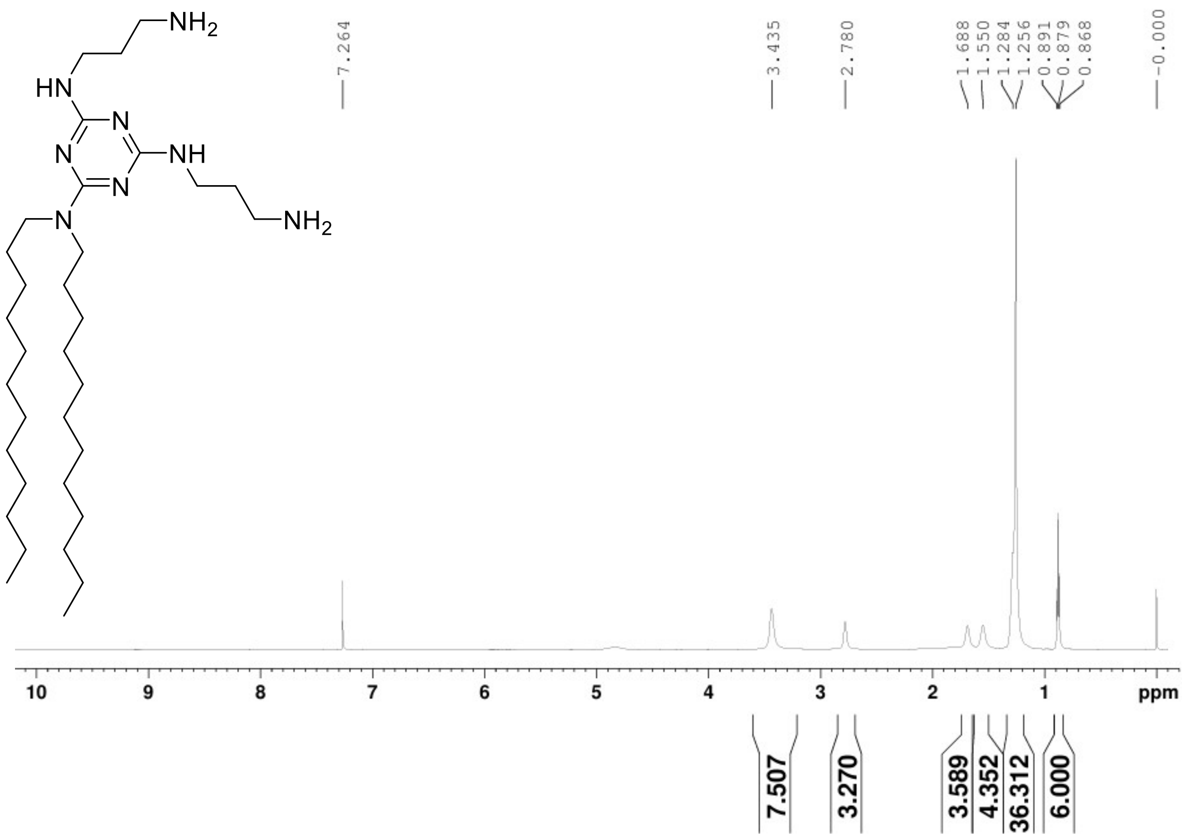


**Figure S8A:** $^{1}H$ NMR Spectra of Lipid TZ C12

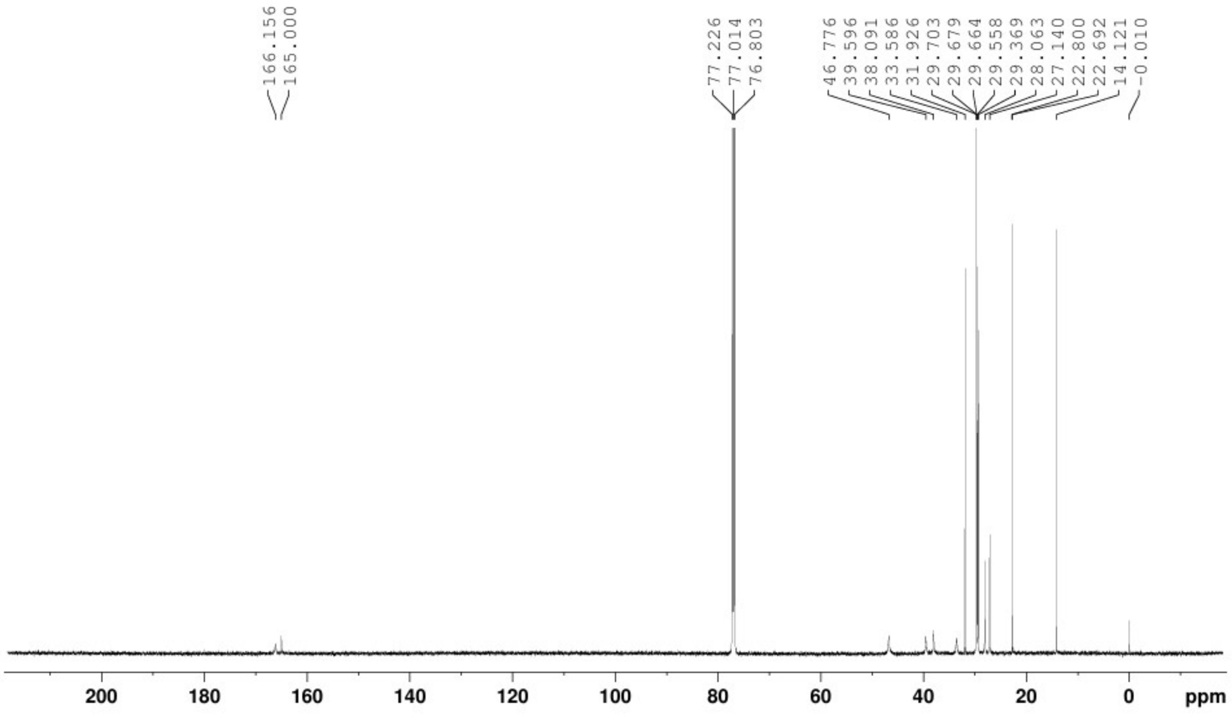


**Figure S8B:** $^{13}C$ NMR Spectra of Lipid TZ C12

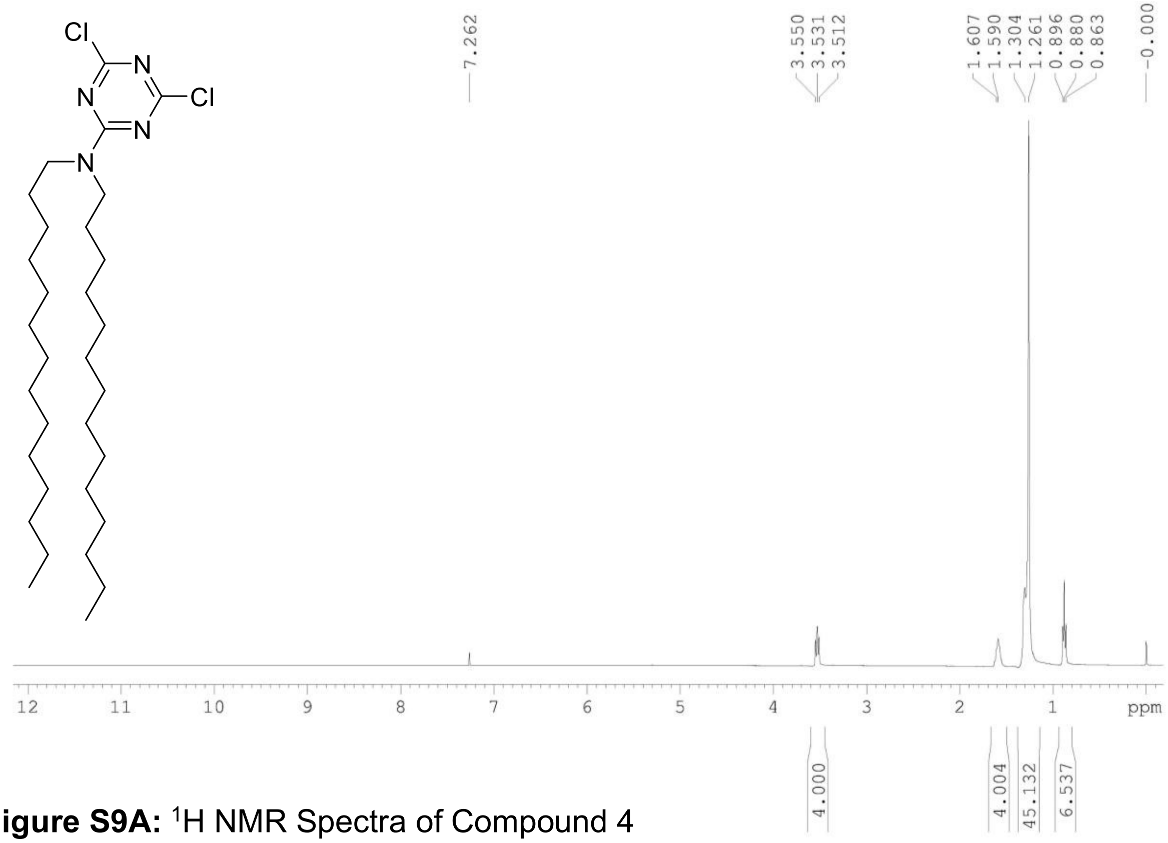


**Figure S9A:** $^{1}$H NMR Spectra of Compound 4

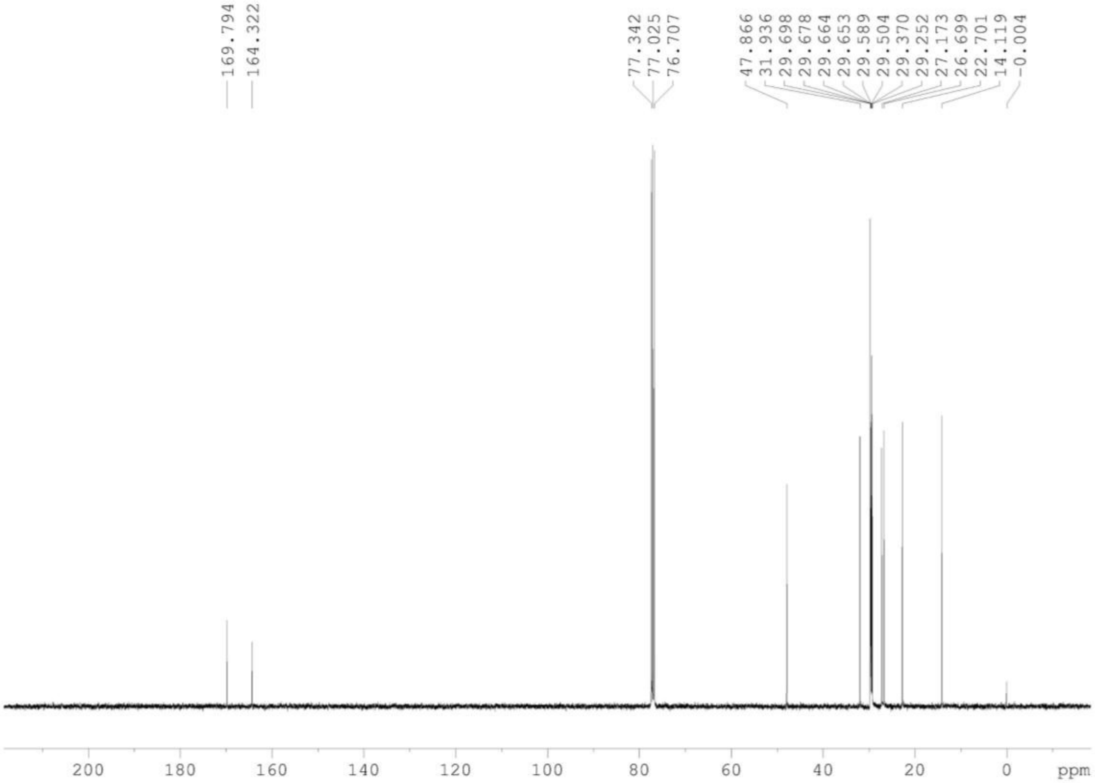


**Figure S9B:** $^{13}$C NMR Spectra of Compound 4

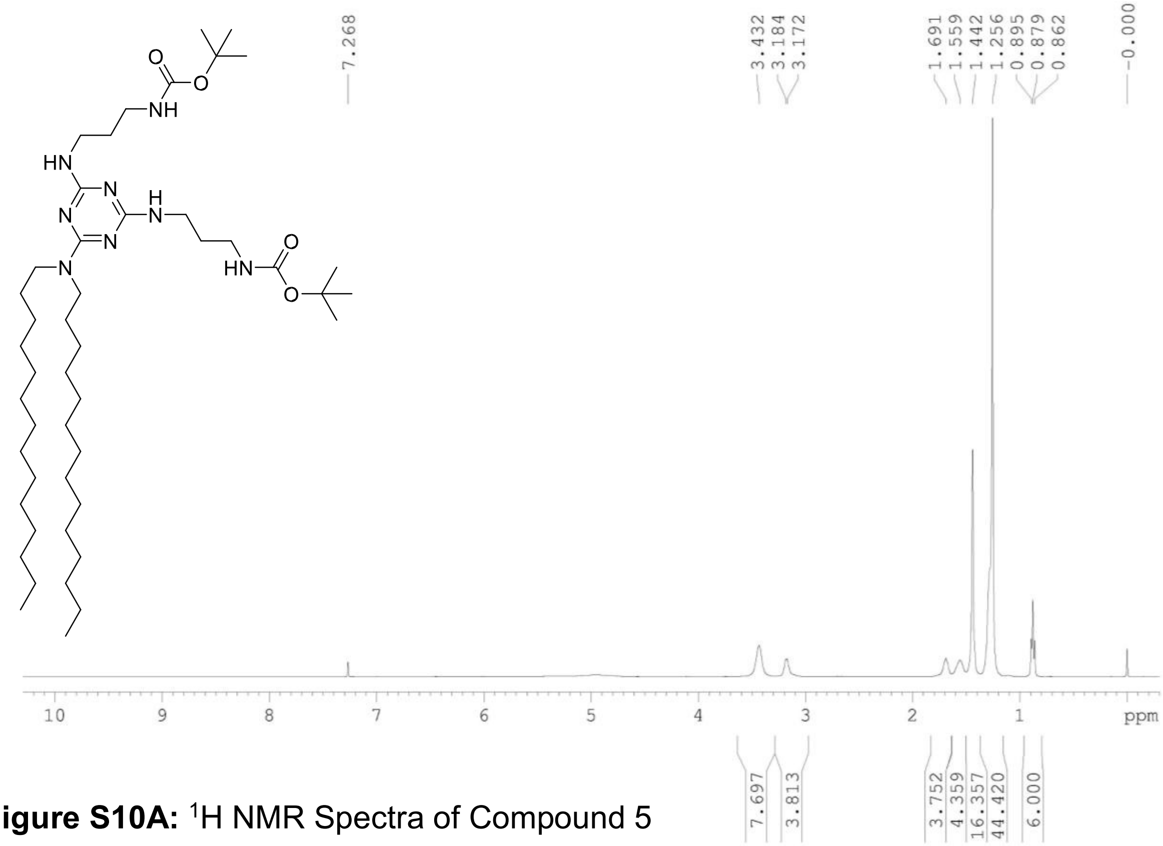


**Figure S10A:** $^{1}H$ NMR Spectra of Compound 5

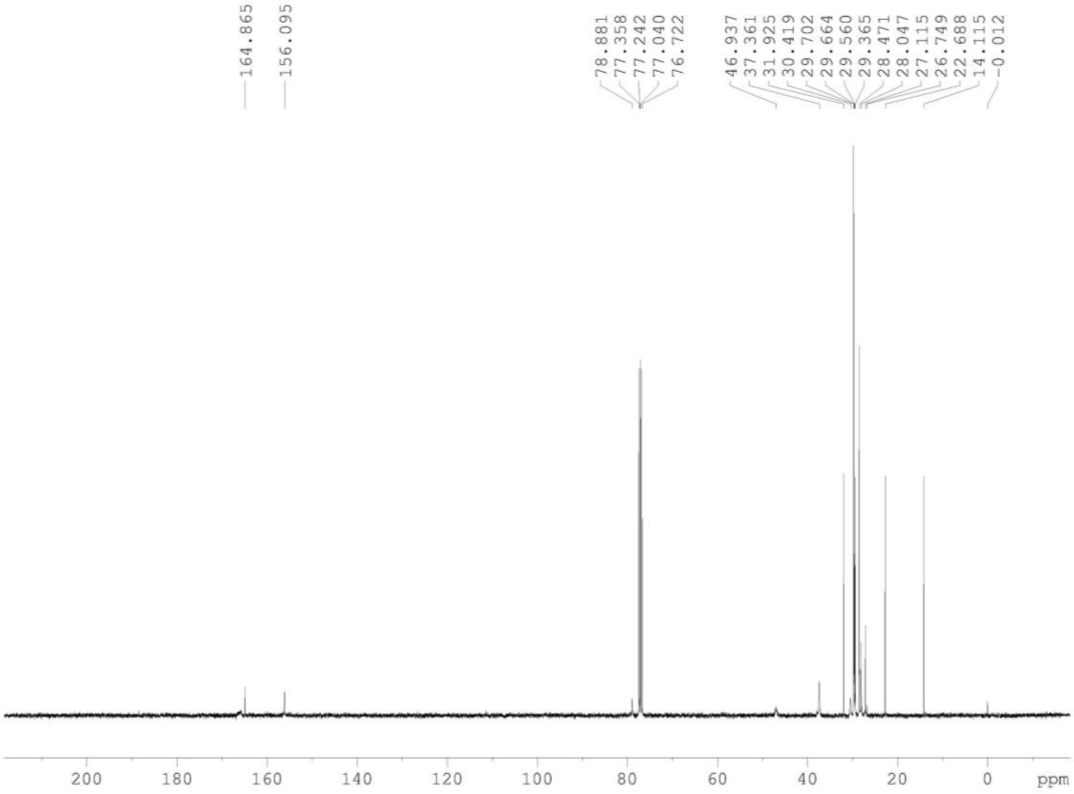


**Figure S10B:** $^{13}C$ NMR Spectra of Compound 5

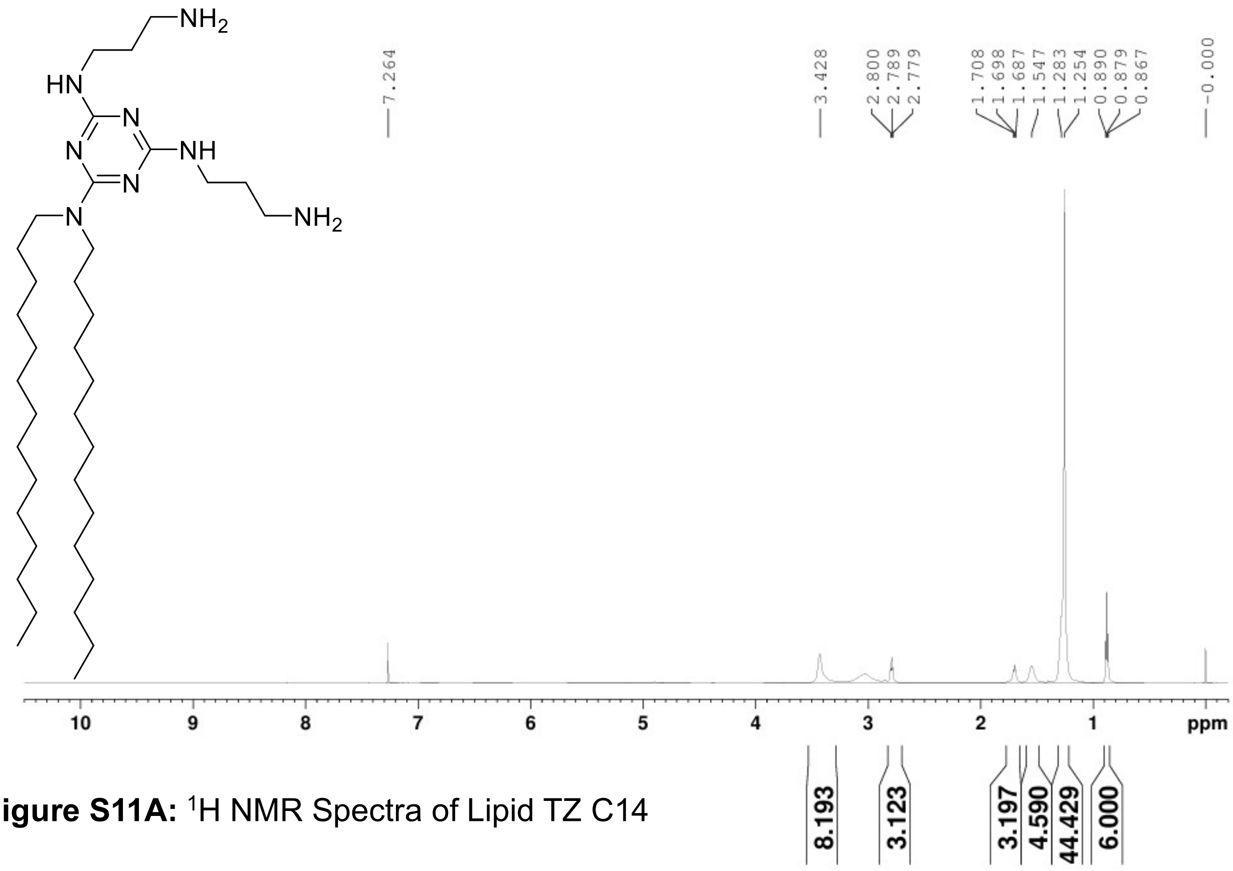

**Figure S11A:** $^{1}H$ NMR Spectra of Lipid TZ C14

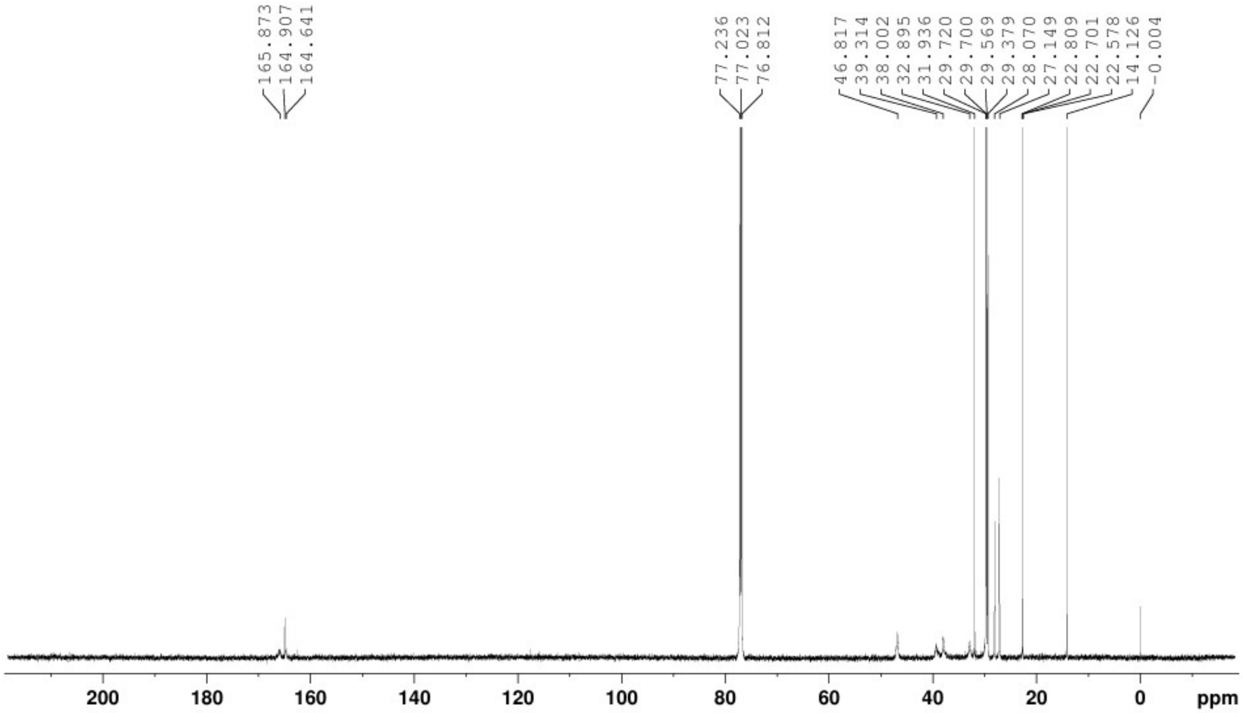

**Figure S11B:** $^{13}C$ NMR Spectra of Lipid TZ C14

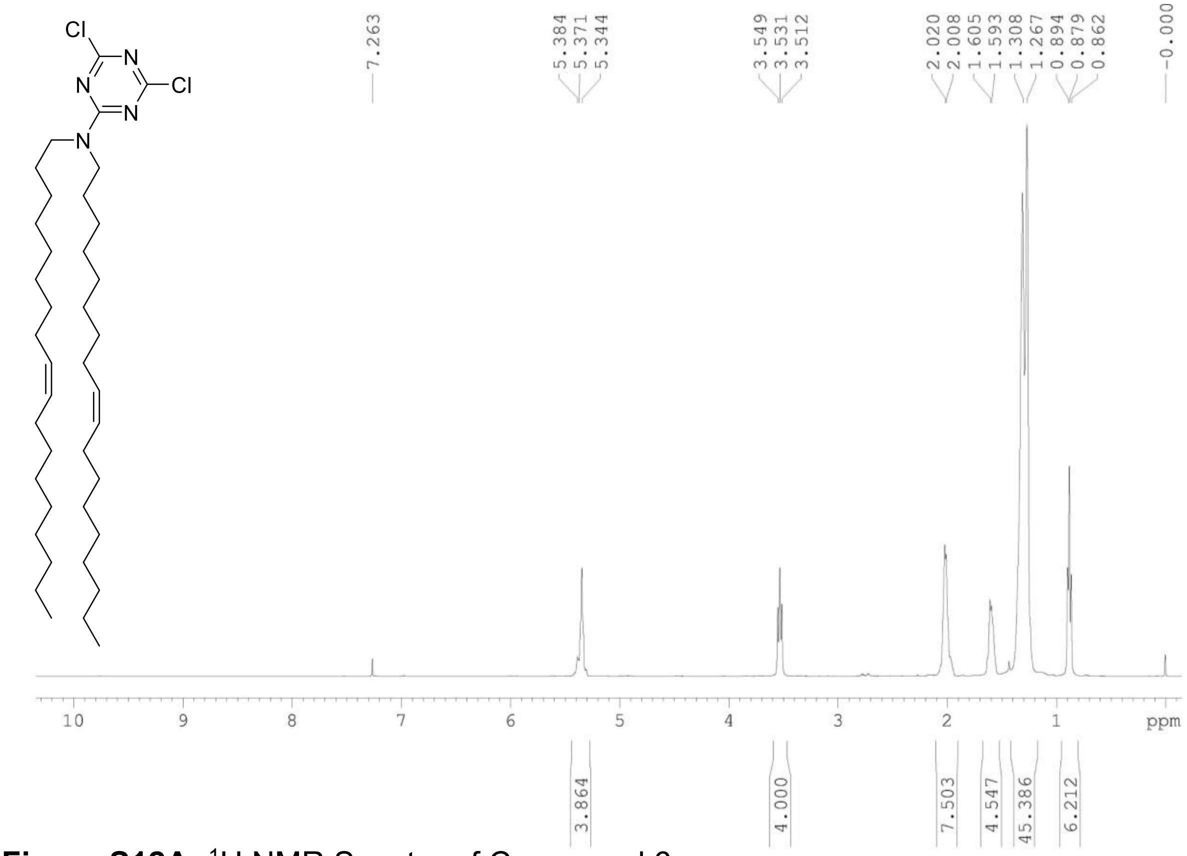


**Figure S12A:** $^{1}$H NMR Spectra of Compound 6

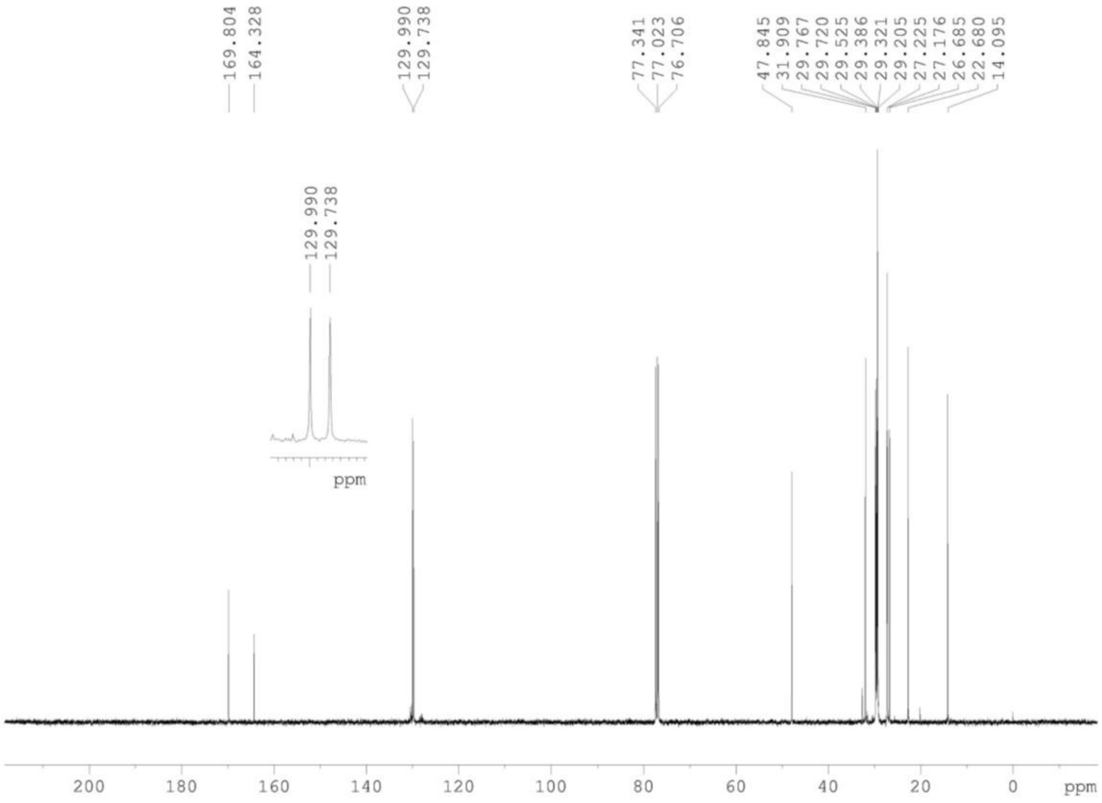


**Figure S12B:** $^{13}$C NMR Spectra of Compound 6

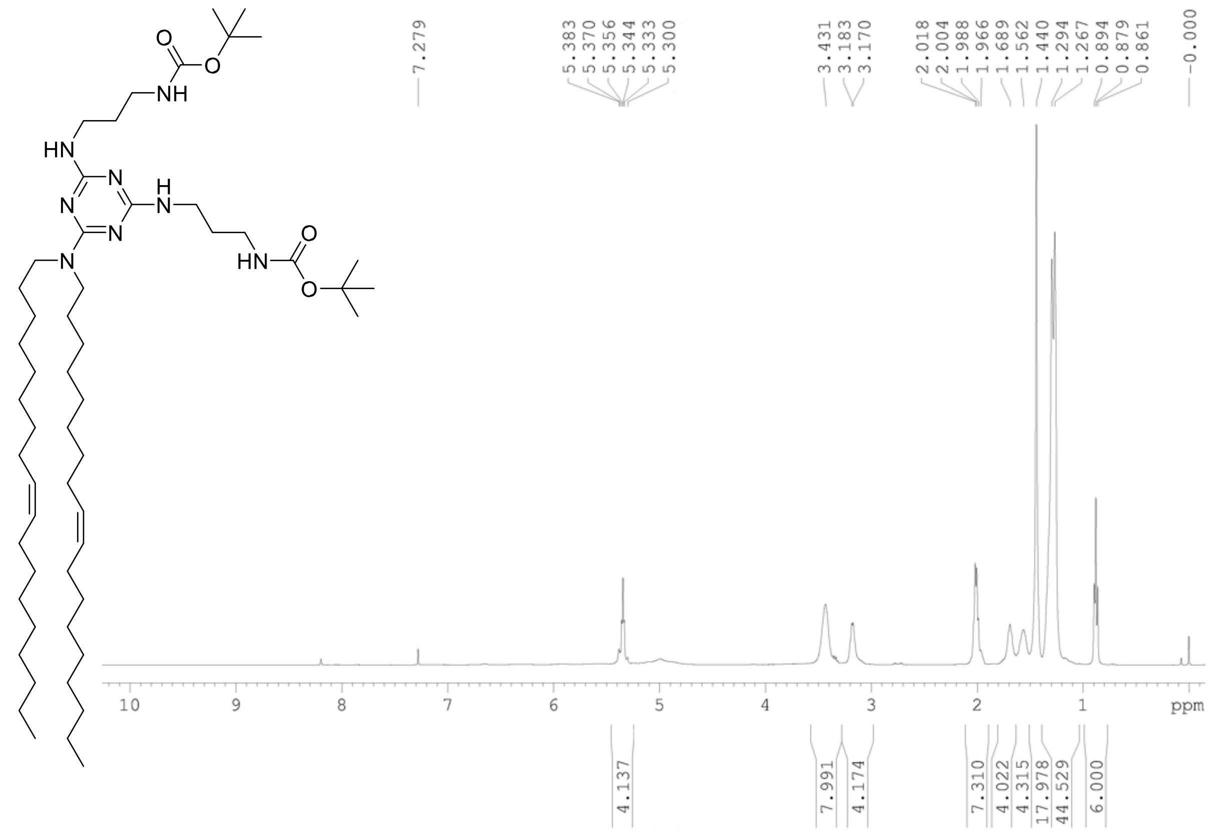


**Figure S13A:** $^{1}$H NMR Spectra of Compound 7

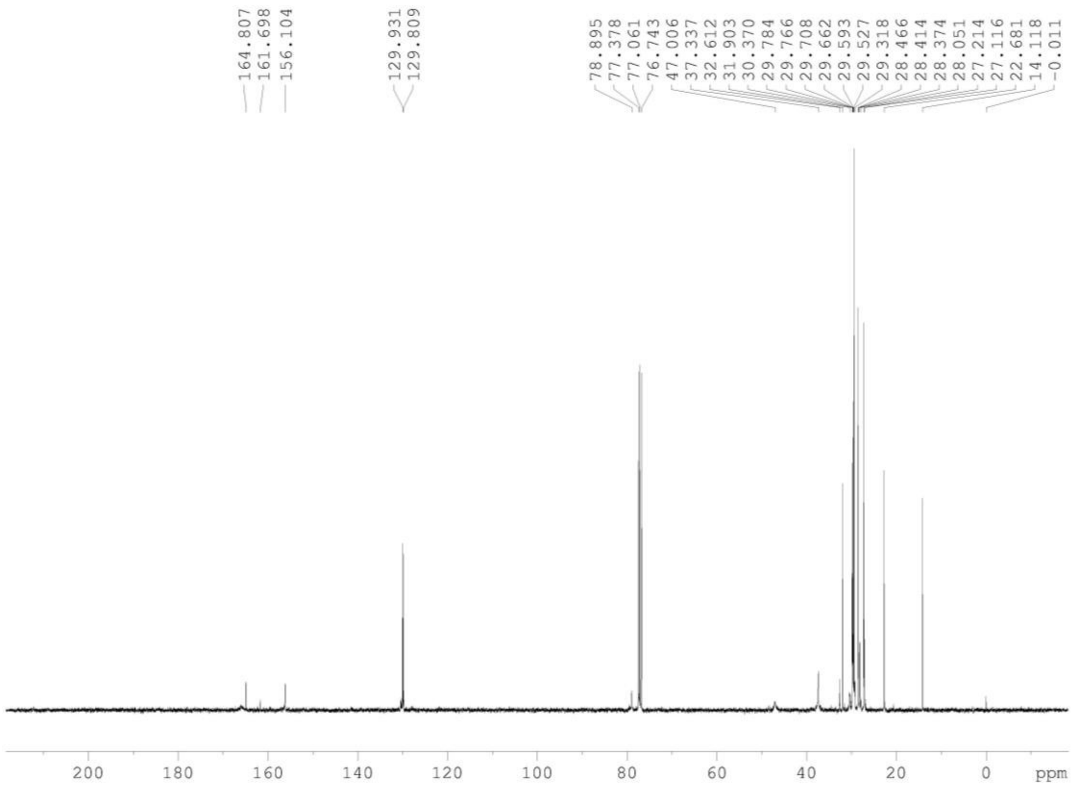


**Figure S13B:** $^{13}$C NMR Spectra of Compound 7

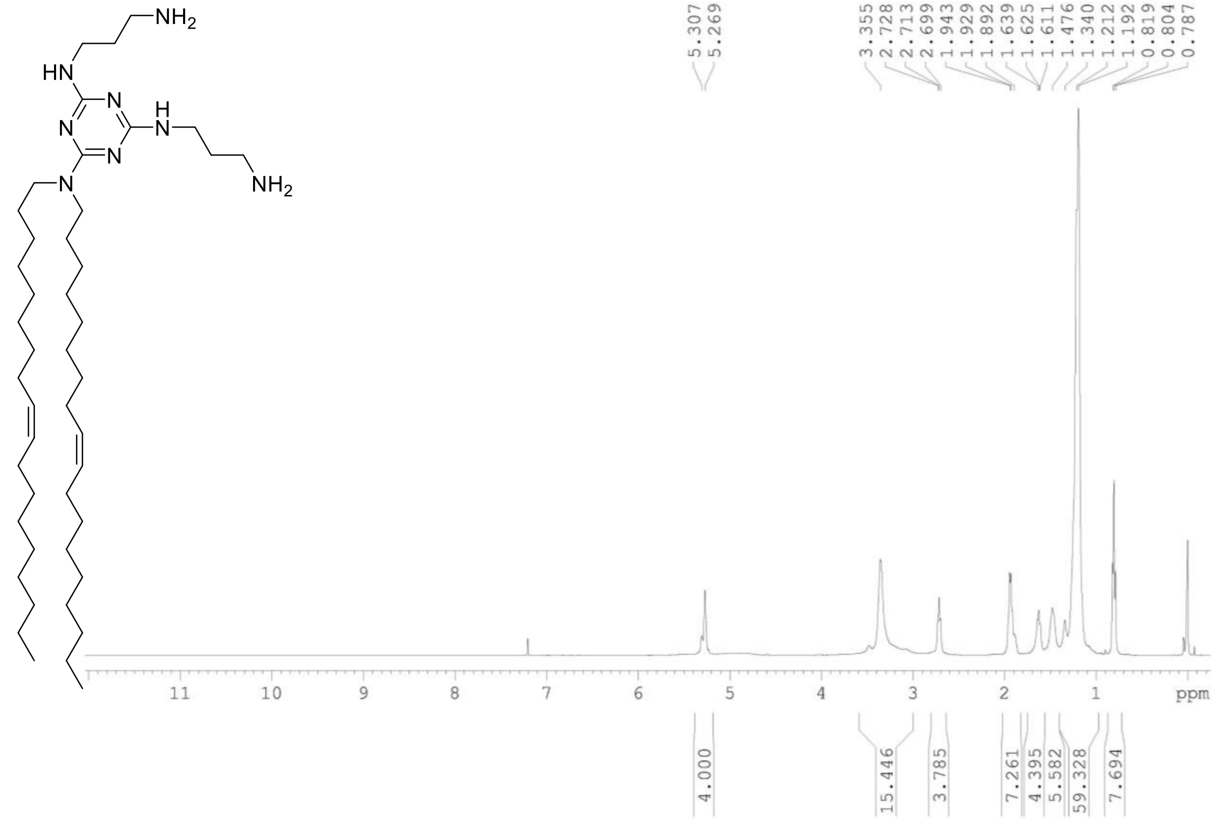


**Figure S14A:** $^{1}$H NMR Spectra of Lipid TZ C18:1

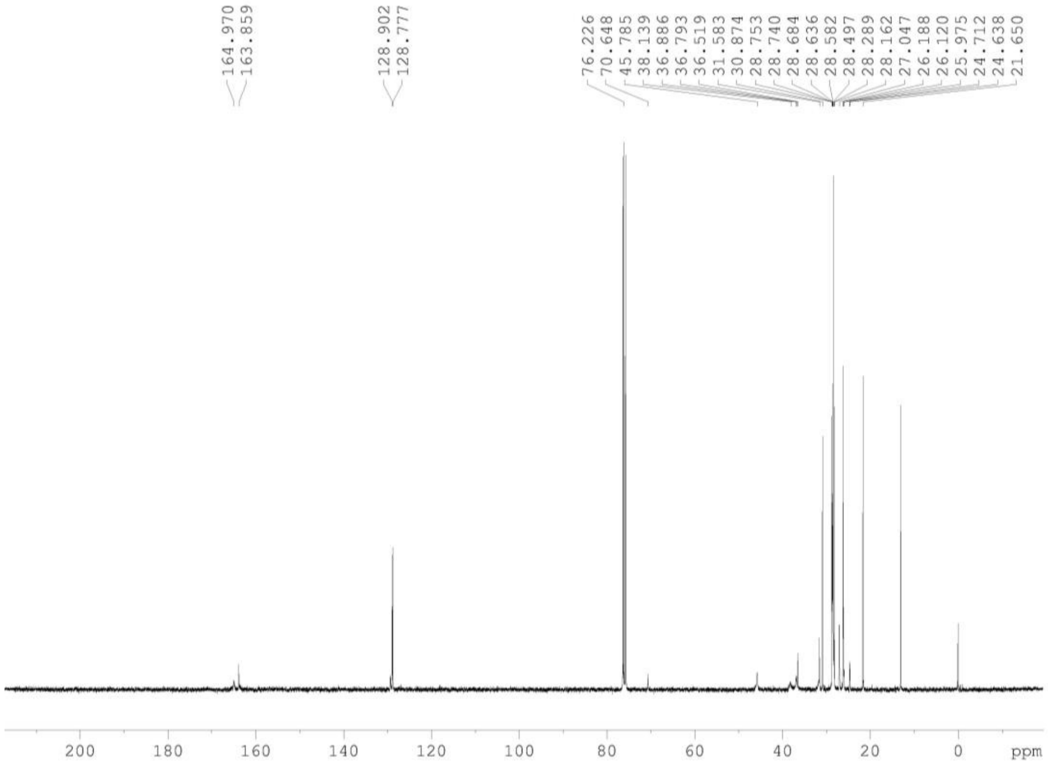


**Figure S14B:** $^{13}$C NMR Spectra of Lipid TZ C18:1

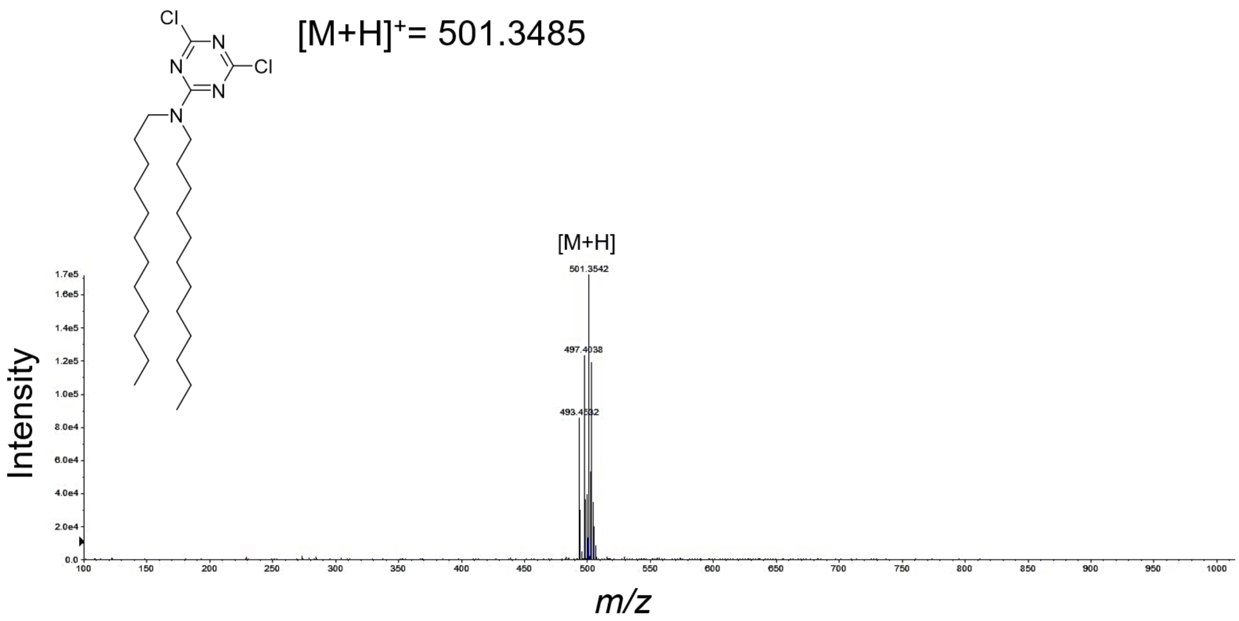


**Figure S15:** HRMS Spectra of Compound 1

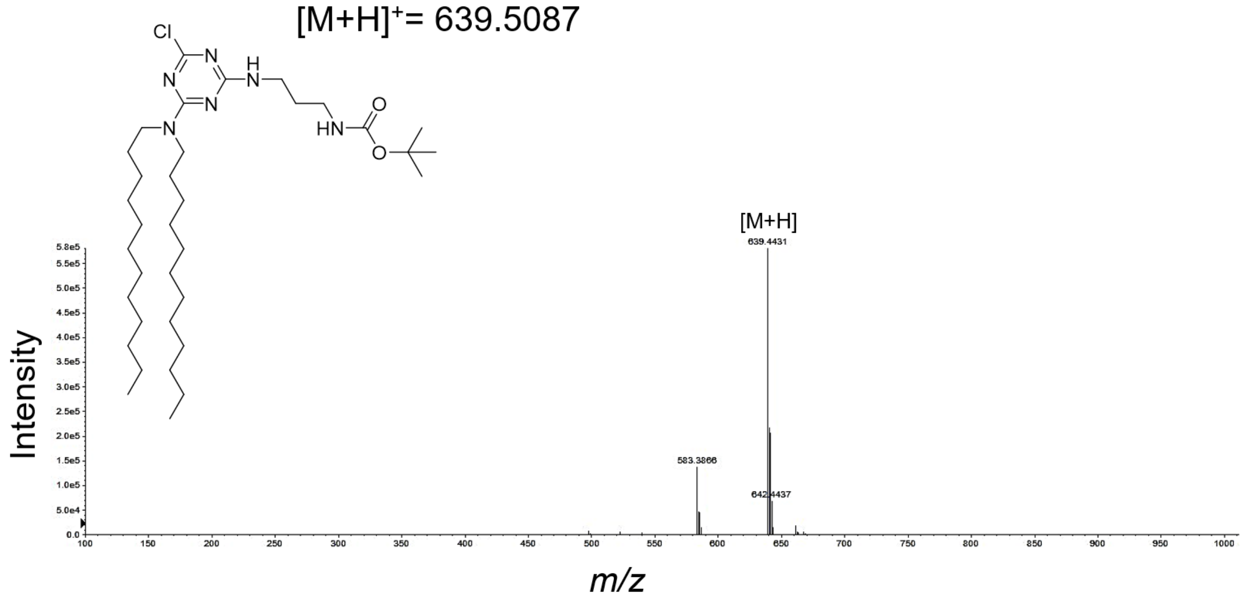


**Figure S16:** HRMS Spectra of Compound 2

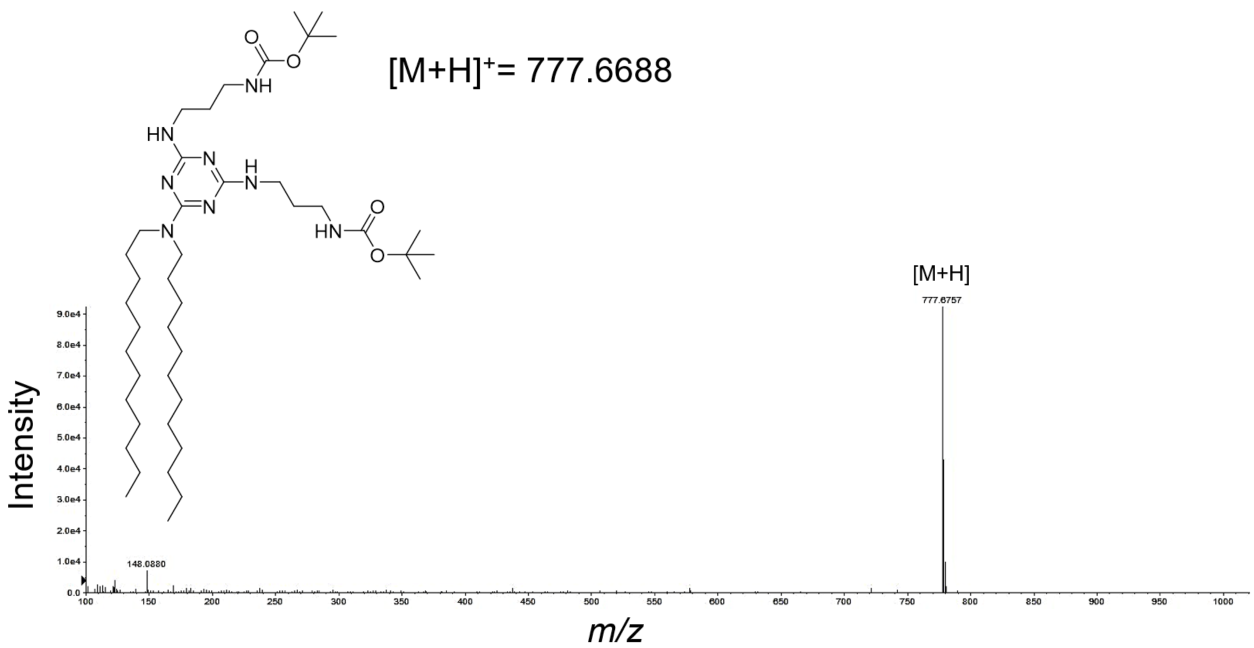


**Figure S17:** HRMS Spectra of Compound 3

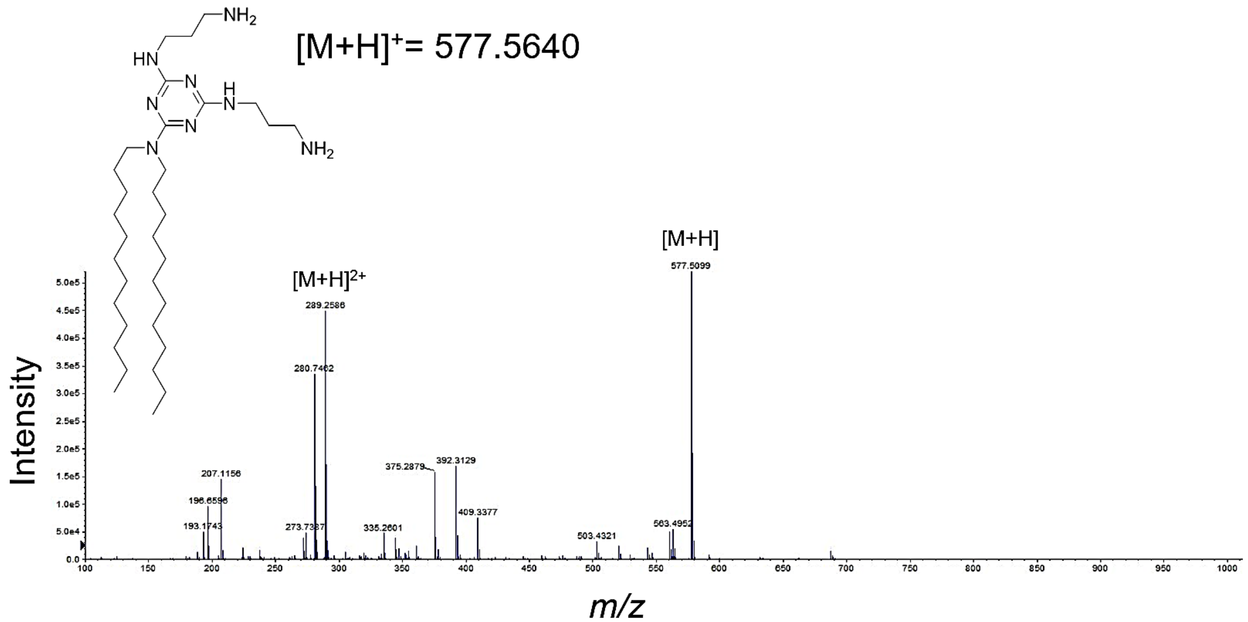


**Figure S18:** HRMS Spectra of Lipid TZ C12

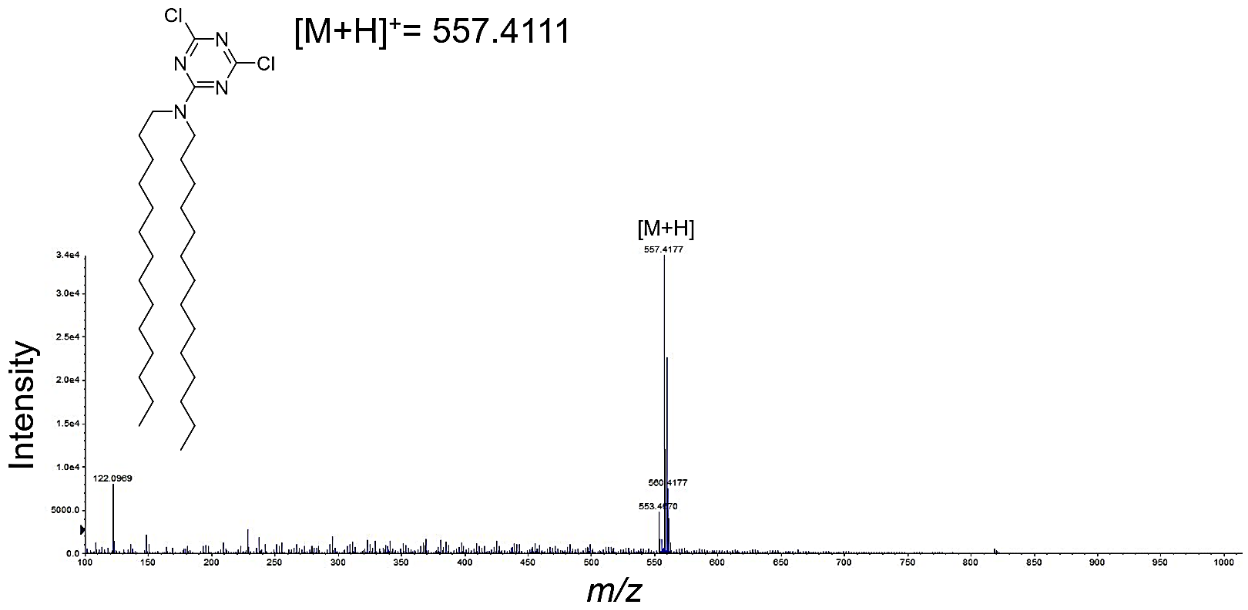


**Figure S19:** HRMS Spectra of Compound 4

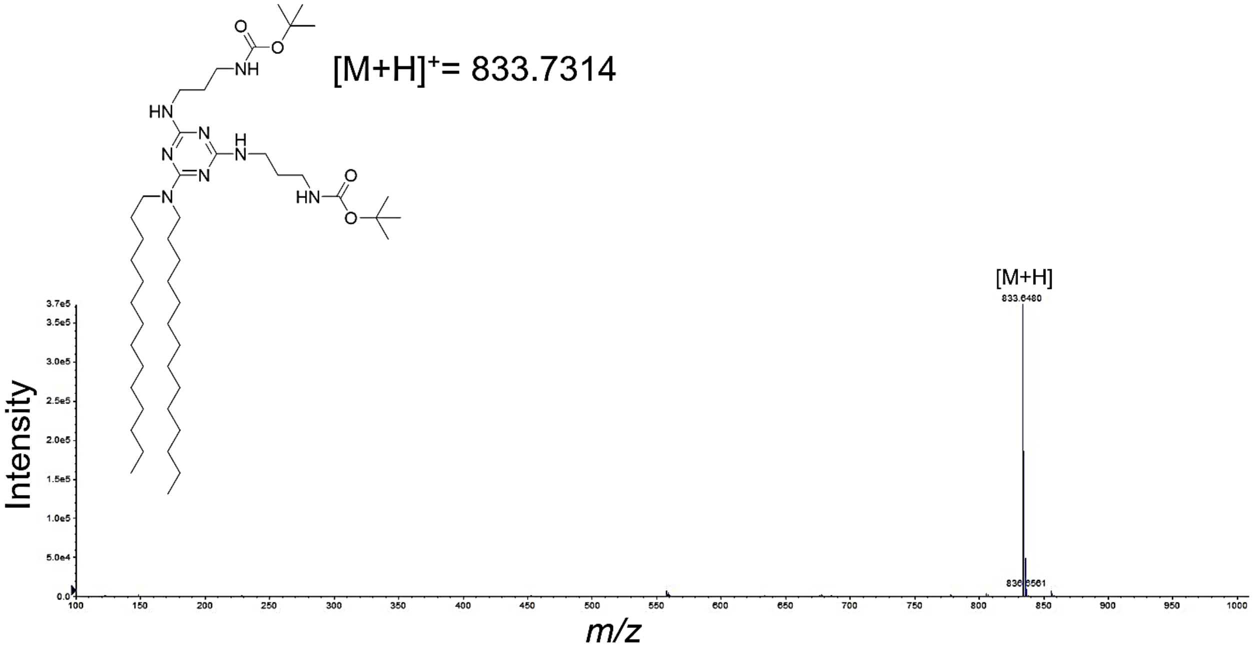


**Figure S20:** HRMS Spectra of Compound 5

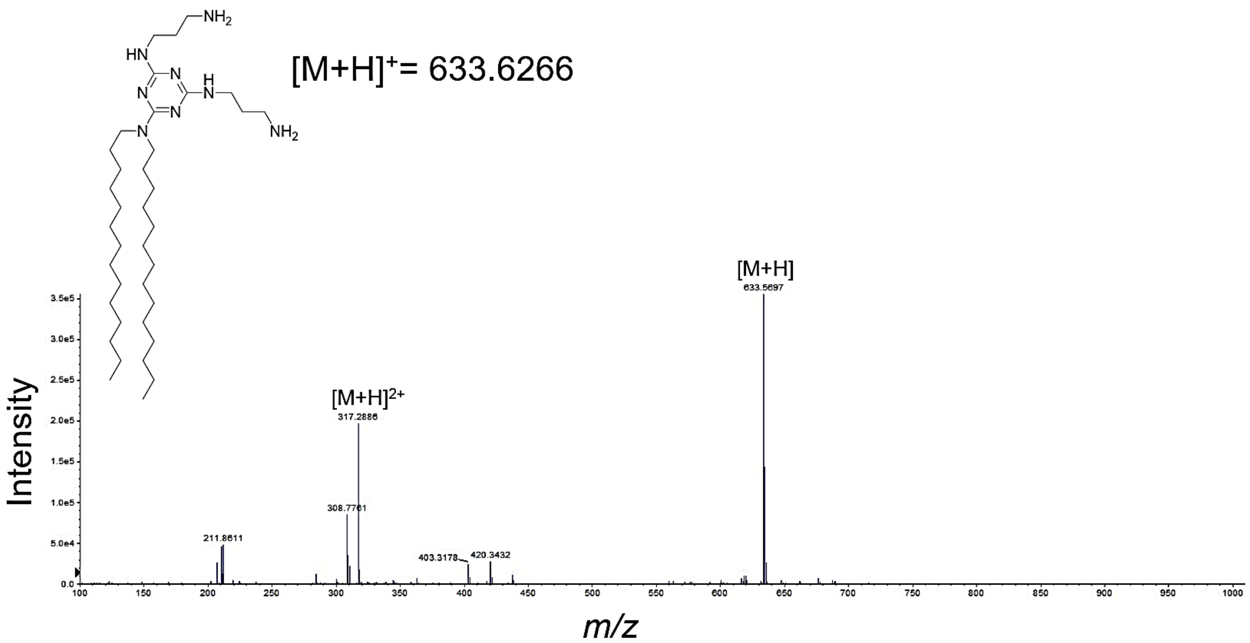


**Figure S21:** HRMS Spectra of Lipid TZ C14

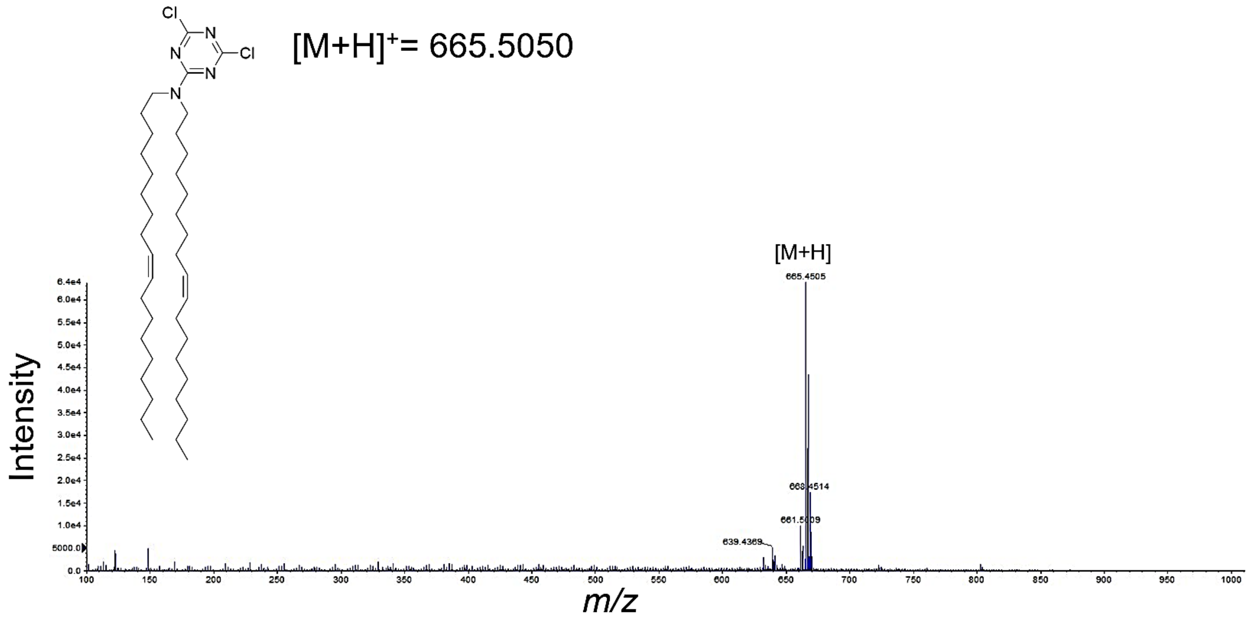


**Figure S22:** HRMS Spectra of Compound 6

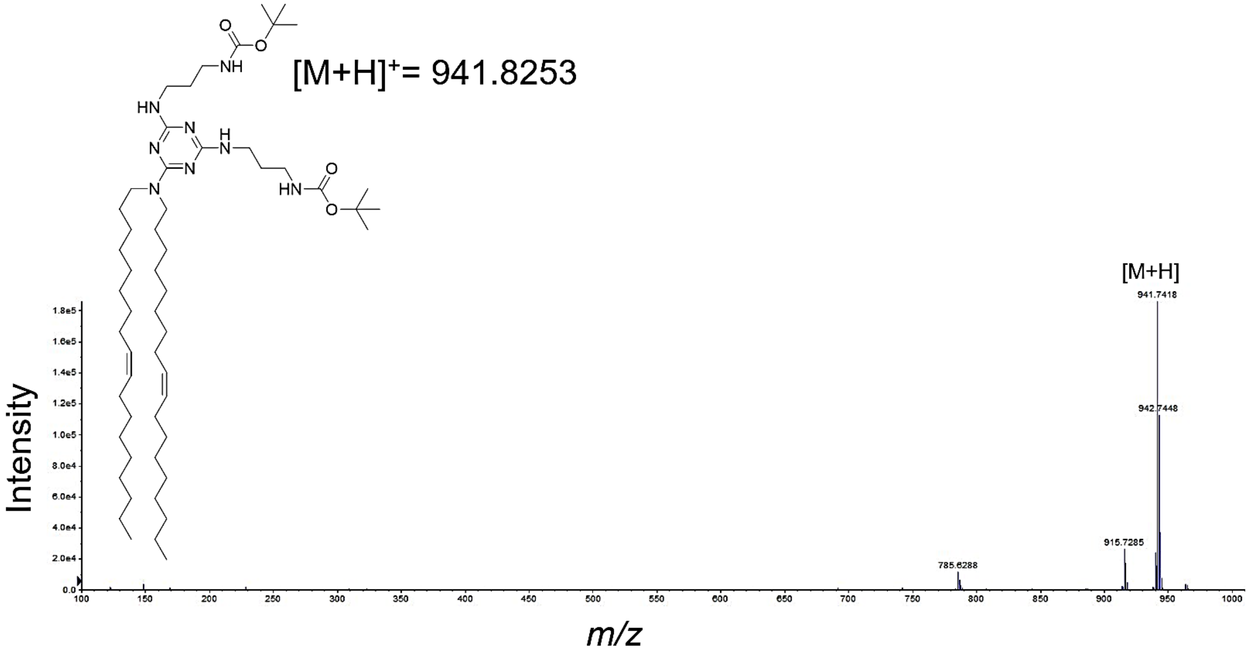


**Figure S23:** HRMS Spectra of Compound 7
